# Cytometry Masked Autoencoder: An Accurate and Interpretable Automated Immunophenotyper

**DOI:** 10.1101/2024.02.13.580114

**Authors:** Jaesik Kim, Matei Ionita, Matthew Lee, Michelle L. McKeague, Ajinkya Pattekar, Mark M. Painter, Joost Wagenaar, Van Truong, Dylan T. Norton, Divij Mathew, Yonghyun Nam, Sokratis A. Apostolidis, Cynthia Clendenin, Patryk Orzechowski, Sang-Hyuk Jung, Jakob Woerner, Caroline A.G. Ittner, Alexandra P. Turner, Mika Esperanza, Thomas G. Dunn, Nilam S. Mangalmurti, John P. Reilly, Nuala J. Meyer, Carolyn S. Calfee, Kathleen D. Liu, Michael A. Matthy, Lamorna Brown Swigart, Ellen L. Burnham, Jeffrey McKeehan, Sheetal Gandotra, Derek W. Russel, Kevin W. Gibbs, Karl W. Thomas, Harsh Barot, Allison R. Greenplate, E. John Wherry, Dokyoon Kim

## Abstract

High-throughput single-cell cytometry data are crucial for understanding involvement of immune system in diseases and responses to treatment. Traditional methods for annotating cytometry data, specifically manual gating and clustering, face challenges in scalability, robustness, and accuracy. In this study, we propose a cytometry masked autoencoder (cyMAE), which offers an automated solution for immunophenotyping tasks including cell type annotation. The cyMAE model is designed to uphold user-defined cell type definitions, thereby facilitating easier interpretation and cross-study comparisons. The cyMAE model operates on a pre-train and fine-tune approach. In the pre-training phase, cyMAE employs *Masked Cytometry Modelling (MCM)* to learn relationships between protein markers in immune cells solely based on protein expression, without relying on prior information such as cell identity and cell type-specific marker proteins. Subsequently, the pre-trained cyMAE is fine-tuned on multiple specialized tasks via task-specific supervised learning. The pre-trained cyMAE addresses the shortcomings of manual gating and clustering methods by providing accurate and interpretable predictions. Through validation across multiple cohorts, we demonstrate that cyMAE effectively identifies co-occurrence patterns of bound labeled antibodies, delivers accurate and interpretable cellular immunophenotyping, and improves the prediction of subject metadata status. Specifically, we evaluated cyMAE for cell type annotation and imputation at the cellular-level and SARS-CoV-2 infection prediction, secondary immune response prediction against COVID-19, and prediction of the infection stage in COVID-19 progression at the subject-level. The introduction of cyMAE marks a significant step forward in immunology research, particularly in large-scale and high-throughput human immune profiling. This approach offers new possibilities for predicting and interpreting cellular-level and subject-level phenotypes in both health and disease.

## Introduction

High-throughput single-cell protein expression data, acquired through flow and mass cytometry, are essential to understanding the role of the immune system in infectious diseases, autoimmunity, cancer, and the response of immune cells post-treatment. Cytometry assays are designed to profile millions of cells from a biological sample, precisely quantifying biomarkers specific to various cell types. In contrast, single-cell RNA sequencing (scRNA-seq) approaches are generally on the scale of thousands of cells. This substantial difference in scale grants cytometry significant advantages for the identification and characterization of rare cell population and enhances the overall comprehensiveness of the data collected. For example, cytometry can pinpoint cell populations that are differentially abundant or proteins that are differentially expressed between subject groups. This process of immune profiling effectively delineates both similarities and diversities within the immune landscape of different subjects, contributing significantly to precision medicine by enabling predictions at an individual level.

The most prevalent approach for analyzing cytometry data is manual gating, a process involving user-applied sequential filters to bivariate plots of protein markers, thereby isolating specific cell subsets for focused analysis^1^. These bivariate plots visually represent the distribution of protein markers, allowing a human analyst to manually identify and select cells based on their prior knowledge of these distributions. Despite widespread use of this approach, manual gating faces several significant challenges. Firstly, it is a time-intensive process, particularly for panels with over a dozen markers^2,3^, as the number of biaxial plots to consider increases quadratically with the number of parameters measured. Secondly, manual gating is prone to subjectivity and bias^2,3^. Each analysis is influenced by pre-existing knowledge, which can lead to a bias towards anticipated results. Subjectivity also enters through the selection of the order of marker combinations and the definition of gate boundaries. Additionally, due to panel size limitation, cytometry panel designs often restrict the search space for pre-defined markers and the corresponding cell types. Thirdly, results from manual gating can be challenging to replicate^2,3^. Different studies may employ varied gating strategies, including distinct gating sequences, shapes, and boundaries for gates, impacting the robustness and consistency of identified cell subsets. Moreover, the level of gating stringency also varies between individual analysts, contributing to inconsistent results.

The ability to simultaneously measure multiple protein markers has significantly increased the complexity of cytometry data. This complexity has led to the development of automated analysis techniques, particularly unsupervised clustering methods like FlowSOM^4^, PhenoGraph^5^, Scaffold Maps^6^, and X-shift^7^. Although these clustering approaches address some limitations of manual gating, they also introduce their own set of constraints. Notably, while unsupervised clustering methods can detect data variability, they struggle to differentiate between biological or technical sources of this variability. This limitation makes these methods susceptible to batch effects, shifts in data distribution, and non-specific binding of antibodies^8^. Another challenge arises in cross-study comparisons, where minor variations in panel selection, sample collection, measurement noise, or random seeding can lead to abrupt changes in cluster boundaries. For example, CD4 T cells might be clustered differently in studies based on memory or functional subtypes, complicating direct comparisons between even highly overlapping datasets. To strike a balance between labor-intensive manual analysis and unpredictable unsupervised analysis, we focus on combining unsupervised and supervised learning to develop an automated method that can immunophenotype future samples using consistent cell type ontologies, regardless of experimental variations.

In the broader context of big data and advanced computational models, artificial intelligence (AI) has achieved great success in fields like computer vision and natural language processing. The effort needed to manually label data makes it extremely difficult to fully leverage the vast amounts of existing unlabeled data in the supervised learning paradigm. However, the revolution of self-supervised learning techniques, particularly in the pre-training phase, empowers models to more accurately learn data distributions and utilize unlabeled data effectively. The core concept behind self-supervised pre-training is randomly masking a portion of the input data and training the model to reconstruct masked information using context clues from the surrounding data. This approach allows the pre-trained model to be fine-tuned for specific downstream tasks or to function as generative AI. Coupling the transformer^9^ architecture, known for its high expressiveness and scalability, has led to significant synergistic effects. Notable examples include *Masked Language Modeling (MLM) as seen in* BERT^10^ and GPT^11^ and *Masked Image Modeling (MIM)* in models like ViT^12^, BeiT^13^, and MAE^14^ in computer vision.

The success of the masking approach has reverberated within the biomedical field as well. For example, protein language models (pLMs) are a set of AI models trained on extensive sets of unlabeled protein sequences^15-17^. pLMs have steadily gained traction across diverse applications for protein design, including antibody engineering^18^ and drug discovery^19,20^. In addition, AI models trained on unlabeled single-cell RNA sequencing (scRNA-seq) data have been published and used for cell annotation purposes^21-27^. Thus, masking models have proven to substantially outperform previous conventional methods in effectiveness and show great potential in biomedical applications. Similarly, we apply these techniques for immunophenotyping, as cytometry data can be structured in a similar way.

In this study, we develop an accurate and interpretable automated immunophenotyper for single-cell cytometry data, using a technique we call *Masked Cytometry Modelling (MCM)*. This approach employs self-supervised pre-training on single-cell cytometry data. During *MCM*, our model learns the relationships and dependencies among markers on immune cells by analyzing expression patterns in the massive amount of data sets, without requiring additional information about cellular identity. The resulting pre-trained model can then export a useful representation that is advantageous for various downstream tasks, surpassing the utility of the original data. We demonstrate that our model not only overcomes the challenges of manual gating and clustering methods but also provides accurate results even on independent datasets that were never seen during its training. This model can accurately identify complex cell types and interprets which crucial protein markers predict targets. Moreover, the model exhibits scalability, reproducibility, and enhanced precision in subject-level phenotyping. In contrast to previous approaches, our model offers a novel approach combining pre-training and fine-tuning for automated annotation of cell identity in single-cell cytometry data. This novel approach in immunophenotyping promises to broaden the impact of existing cytometry data and enhance immunological knowledge by more accurately phenotypes at both cellular and subject levels.

## Results

### Cytometry Masked Autoencoder (cyMAE) algorithm

To address the challenges of time-consuming and labor-intensive immunophenotyping in cytometry data, we propose cyMAE, a cytometry Masked Autoencoder model. This innovative model constructs and employs latent embeddings of single-cell cytometry data to obtain state-of-the-art performance on various cell-level and subject-level tasks. cyMAE is built upon a Masked Autoencoder (MAE)^14^ architecture, featuring stacked transformer blocks in both the encoder and decoder. Inspired by successful methodologies in computer vision and natural language processing, cyMAE undergoes a two-phase training process: self-supervised pre-training followed by supervised fine-tuning, as illustrated in **Figure 1a**. The main advantage of this approach is its ability to use large-scale, easily obtainable unlabeled data during the initial self-supervised pre-training phase, thus reducing the reliance on scarce and labor-intensive labeled data in the subsequent fine-tuning phase. During pre-training, a randomly selected subset of the protein expression data is masked and fed to an encoder, which produces latent embeddings of the masked data. In turn, these embeddings are processed by a decoder that attempts to reconstruct the unmasked, original data (**Figure 1b, Supplementary Figure 1**). Through this process, the encoder-decoder system learns to optimize the embeddings to minimize reconstruction error, effectively enabling the model to obtain informative data embeddings without requiring explicit ground truth labels. During the second fine-tuning stage, the model employs the full, unmasked protein expression data to generate latent cell representations using the encoder that was pre-trained in the first stage. These representations are applicable to a range of downstream tasks, whether they involve labeled data or not. Cell representations generated by the pre-trained encoder can be used for unsupervised tasks or plugged into another classifier to solve tasks through supervised fine-tuning. Specifically, we evaluated the pre-trained cyMAE’s performance on two cell-level tasks: cell type annotation and imputation. Moreover, for subject-level tasks, we tested SARS-CoV-2 infection prediction, secondary immune response prediction against COVID-19, and prediction the infection stage in the COVID-19 progression.

**Figure 1.**
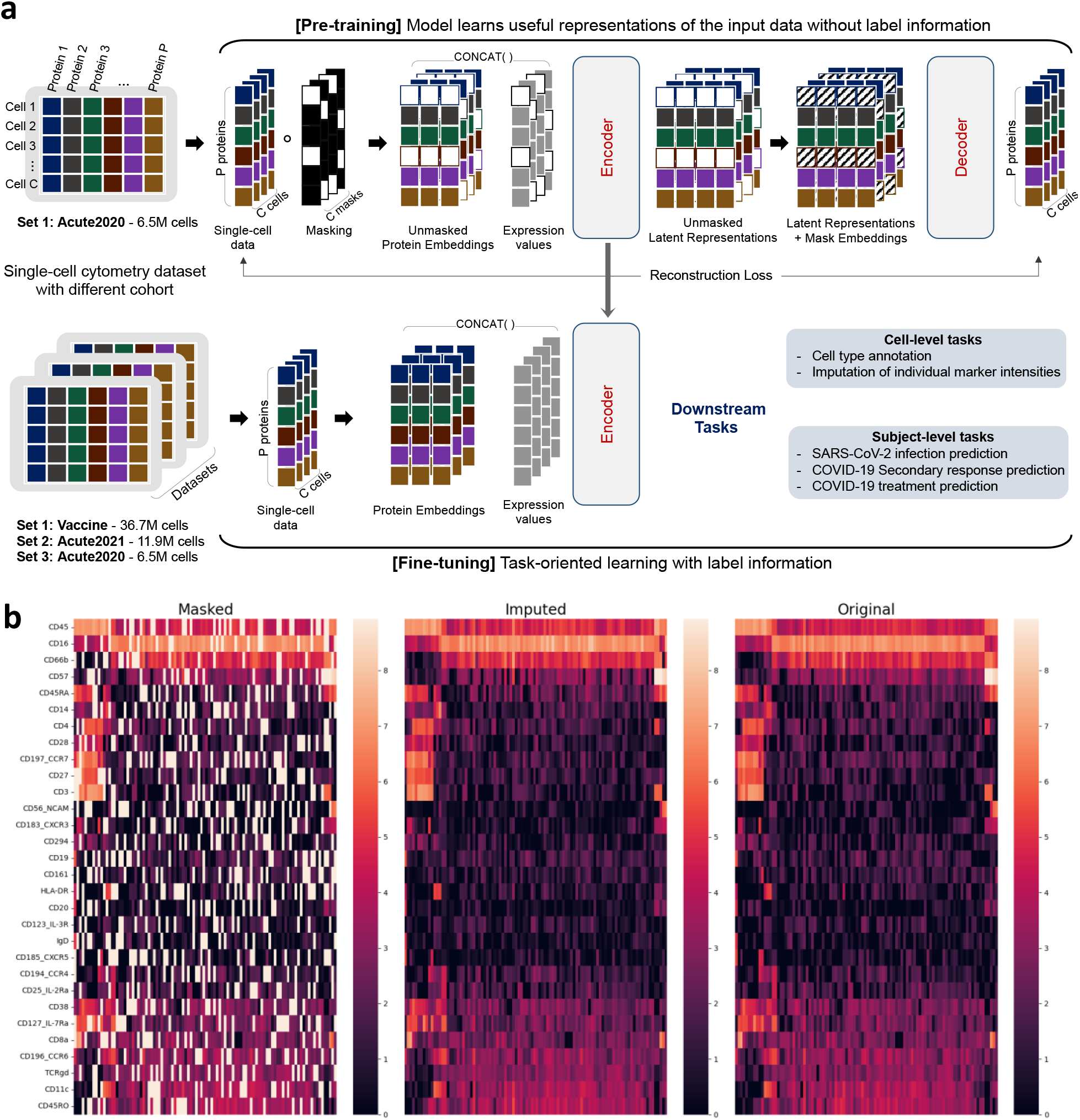
(a) Overview of Cytometry Masked Autoencoder (cyMAE). In the pre-training step, protein expression data is randomly masked. The unmasked protein expressions are concatenated with learnable protein embeddings and fed into the encoder. This encoder generates unmasked latent representations, which are merged with learnable mask embeddings and fed to the decoder for reconstruction of the masked values. In the fine-tuning step, the pre-trained encoder produces latent representations for both cells and subjects, facilitating cell-level and subject-level downstream tasks, respectively. (b) From left to right, masked, imputed (reconstructed), and original data. Each row represents a marker protein, and each column represents a randomly sampled cell. Initially, 25% of the original data is randomly masked, shown in white in the masked data visualization. cyMAE effectively reconstructs these masked regions, demonstrating the model’s accuracy.

We analyzed Cytometry by Time Of Flight (CyTOF) data from three distinct COVID-19 studies conducted at the University of Pennsylvania, referred to as the Acute2020 dataset, Vaccine dataset, and Acute2021 dataset. For all datasets, whole blood was stained with a 30-marker panel. Each of the datasets underwent a routine manual gating practice executed by domain experts to extract single, intact cells in preparation for downstream analysis. The Acute2020 dataset consists of single-time-point samples from 13 patients hospitalized with acute SARS-CoV-2 infection in 2020 and 13 healthy controls, comprising a total of 6.5M cells. The Vaccine dataset includes 37 healthy adults followed longitudinally before and after (7 days after second dose) SARS-CoV-2 mRNA vaccine, for a total of 150 FCS files. This dataset is composed of 36.7M cells. Lastly, the Acute2021 dataset contains longitudinal samples from 42 SARS-CoV-2 infected individuals who were enrolled in the I-SPY COVID-19 Trial^28^ in 2021. Samples were collected at the time of hospital admission and 7 days later. This dataset includes 11.9M cells from 56 FCS files. Prior analysis of flow cytometry data from the Acute2020 dataset revealed heterogenous peripheral blood profiles among patients hospitalized with SARS-CoV-2, capturing both common and uncommon cells and cell phenotypes compared to healthy individuals^29^. Thus, the Acute2020 dataset was chosen for pre-training, while all three datasets were used in the downstream evaluations.

### cyMAE learns antibody co-occurrence patterns

The pre-trained cyMAE effectively learns the patterns of co-occurrence among antibodies targeting specific proteins, purely from data, without relying on any prior knowledge. This capability is demonstrated by the way cyMAE groups proteins based on their co-localization on particular cell types, as seen in **Figure 2a**. For example, proteins that appear mostly on T cells (CD3, CD4, CD8, CD28, CD183 etc.) cluster together, as do proteins that mostly appear on B cells (CD19, CD20, CD185, CD196, IgD). Interestingly, TCRgd and CD197 cluster with neutrophil markers (CD16 and CD66b), despite not traditionally being associated with neutrophils. An in-depth analysis showed that anti-TCRgd and anti-CD197 antibodies in this panel were indeed measured in neutrophils, in lower amounts compared to T cells (**Supplementary Figure 2**). In the case of TCRgd, this is likely non-specific binding, whereas for CD197 the reasons could be technical or biological. Either way, because neutrophils are the most abundant cell type, they are the dominant target for background expression of TCRgd and CD197. This result illustrates that cyMAE can effectively capture the contextual relationships and dependencies between immune cell marker expression levels. Thus, this model can be learnable for data patterns, enabling it to successfully perform subsequent downstream tasks.

**Figure 2.**
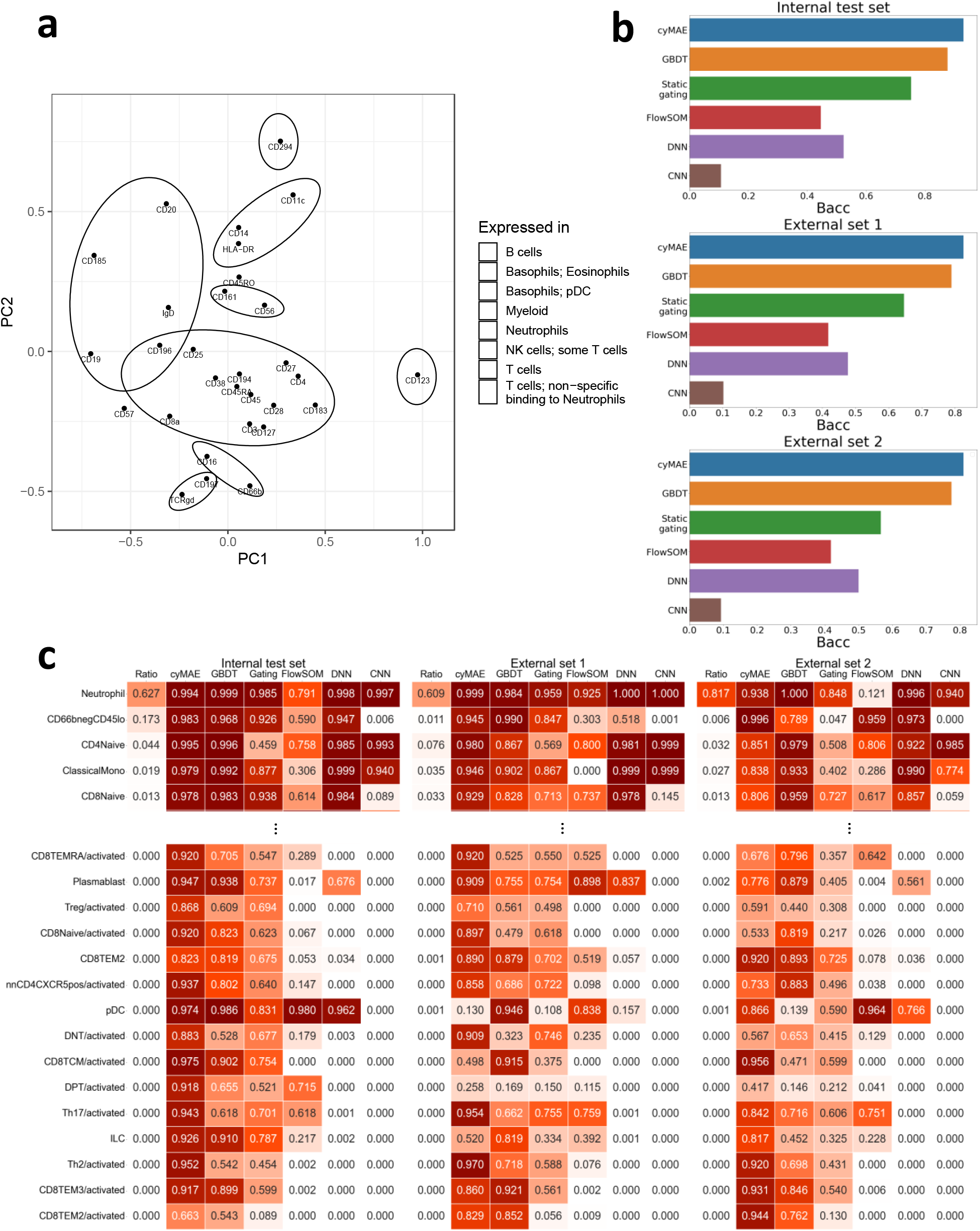
(a) PCA plot of the cyMAE protein embeddings, demonstrating how the model, through unsupervised pre-training, effectively learns protein embeddings that represent the spatial closeness of antibody probes. (b) Model comparisons in the 46 cell type annotation with Balanced Accuracy (Bacc). The internal test set is Vaccine dataset after train-test split, the external set 1 is Acute2021, and the external set 2 is Acute2020. GBDT is a gradient boosting decision tree. Static gating is a method to aggregate into a single consensus gate for each gate in the hierarchy (see Methods). Deep neural network (DNN) denotes the fully-connected neural network proposed by Cheng, L. *et al*.^30^ and Li, H *et al*.^31^ for cytometry data analysis. Convolution neural network (CNN) denotes a model architecture that removes only pooling layer from the CNN proposed by Hu. Z *et al*.^8^ for cytomegalovirus (CMV) classification. (c) Accuracy of cell type annotation for both 5 abundant and 15 rare cell types.

### cyMAE is an accurate cell immunophenotyper

Cell type annotation, traditionally achieved through manual gating and clustering methods, is now efficiently automated by our cyMAE model. By fine-tuning with cell type labels, cyMAE accurately annotates cell types in single-cell datasets. Ground truth labels for 46 cell types, obtained from manual gating, were used (**Supplementary Figure 3**). We used 60% of the Vaccine dataset for fine-tuning the model, 20% as validation and the remaining 20% as an internal test set. We further evaluated cyMAE using the Acute2021 dataset and the Acute2020 dataset as external validation sets (External set 1 and 2, respectively). We compared cyMAE with a gradient boosting decision tree (GBDT)^32^, a fully connected deep neural network (DNN), and a convolutional neural network (CNN) (see Methods) as well as cytometry-specific analysis methods: static gating and unsupervised clustering with FlowSOM.

As a baseline, we took the gating strategy developed on the training dataset and apply it statically to the testing datasets, without adjustments for inter-sample variability (**Supplementary Figure 4**). This approach is equivalent to manually constructing a decision tree and then applying it on the testing data. The other supervised models used here can be seen as refinements of this idea: they attempt to learn a more robust encoding of the gating information by using multivariate rather than bivariate expression patterns. Alongside the supervised classification methods, we included FlowSOM, a popular unsupervised clustering method for cytometry. To match our supervised paradigm, we add an inference mode to FlowSOM by mapping each unseen test datapoint to the nearest SOM node (see Methods).

Given the imbalanced distribution of cell type, with neutrophils comprising over 60% of cells, we used Balanced accuracy (Bacc) to assess model performance fairly. The experimental results showed consistently high Bacc on both internal test sets and two external sets, with the internal test set achieving 93.1% Bacc and the external sets 82.5% and 81.0% Bacc, respectively (**Figure 2b**). When we examined performance by cell type, our model was found to be more accurate than others for most cell types (**Supplementary Figure 5**).

Notably, the model performed particularly well on rare cell types. Accurate prediction of rare cell types is difficult because it is easy for a model to be trained with a bias toward more frequent cell types. However, when comparing performance on cells with a frequency of less than 0.1% in **Figure 2c**, both internal test set and external sets show more accurate predictions for rare cell types than the comparison models in most cases.

In addition, cyMAE’s performance benefits from pre-training, outperforming the cyMAE model from scratch (non-pre-trained), demonstrating the value of leveraging large-scale unlabeled data for pre-training (**Supplementary Table 1**). While FlowSOM scored lower on our accuracy metrics, this outcome does not reflect the quality of the algorithm; instead, it underscores the limitations of unsupervised methods that do not use training labels. This observation illustrates one key pitfall of unsupervised analysis: this approach reveals true variability in the data, which may not be biologically important. For example, unsupervised analysis splits neutrophils, the dominant population, into 6 clusters based on non-specific binding of anti-CD3 or anti-TCRgd, while more subtle, but biologically meaningful, populations like T cell effector memory subsets are missed.

These results show that our cyMAE model is robust to technical variation between datasets, showcasing superior performance across different collection and processing protocols. For example, despite being trained on the Vaccine dataset from cryopreserved samples of healthy subjects in 2021, cyMAE outperformed all other methods on the Acute2020 dataset, which comprised fresh samples from subjects with acute COVID in 2020. These results underline cyMAE’s potential as reliable tool for cell immunophenotyping across diverse datasets.

### cyMAE enhances regression imputation for cytometry data

Current technology for flow and mass cytometry only allows a few dozen markers, and sometimes cost considerations may reduce the number even further. This limitation is unlike single-cell RNA sequencing or other similar single-cell techniques, which can capture thousands of parameters. Despite these limitations, cytometry remains a powerful tool in single-cell biology due to its widespread use, ease of application, clinical implementation, and its capacity to analyze significantly more cells (typically millions versus thousands in single-cell genomics). This latter point means that cytometric approaches are much more robust for interrogating rare cell types often sparsely sampled or missed altogether by single cell sequencing approaches. Fully exploiting these advantages of cytometric approaches through advanced computational methods, for example, allowing measurements on small panel sizes to yield analytical results similar to those on larger panel sizes would be a major advance for the field. To investigate this feasibility, Becht, E. *et al*.^33^ proposed Infinity Flow, applying a Gradient Boosting tree model^32^ to impute the expression of over 300 markers from merely 15. We assessed whether the cyMAE’s cell latent representations could further enhance regression imputation. In our experiments, we masked 7 markers associated with memory subsets in T cells (CD27, CD28, CD45RA, CD45RO, CD127, CD197), using the remaining data to predict the masked marker expressions with both Infinity Flow and cyMAE, the latter fine-tuned for imputation. We used the Acute2020 data for training, and the Vaccine dataset and Acute2021 dataset as external sets (External set 1 and 2, respectively). R-squared values were used for evaluation.

The cyMAE model achieved imputation performances with R-squared values ranging from 0.2-0.6 (**Figure 3a**), despite being limited to 23 markers not directly indicative of T cell memory states and their associated masked markers. Notably, cyMAE outperformed Infinity Flow for five of the seven markers. Beyond identifying patterns of universally expressed proteins, such as CD45RA in NK cells and CD45RO in neutrophils, cyMAE also showed high correlations between true and predicted values, specifically within T cells or for CD27 expression in B cells (**Figure 3b, Supplementary Figure 6-8**). These results suggest that cyMAE can infer information about the memory states of T and B cells, even in the absence of the standard memory markers.

**Figure 3.**
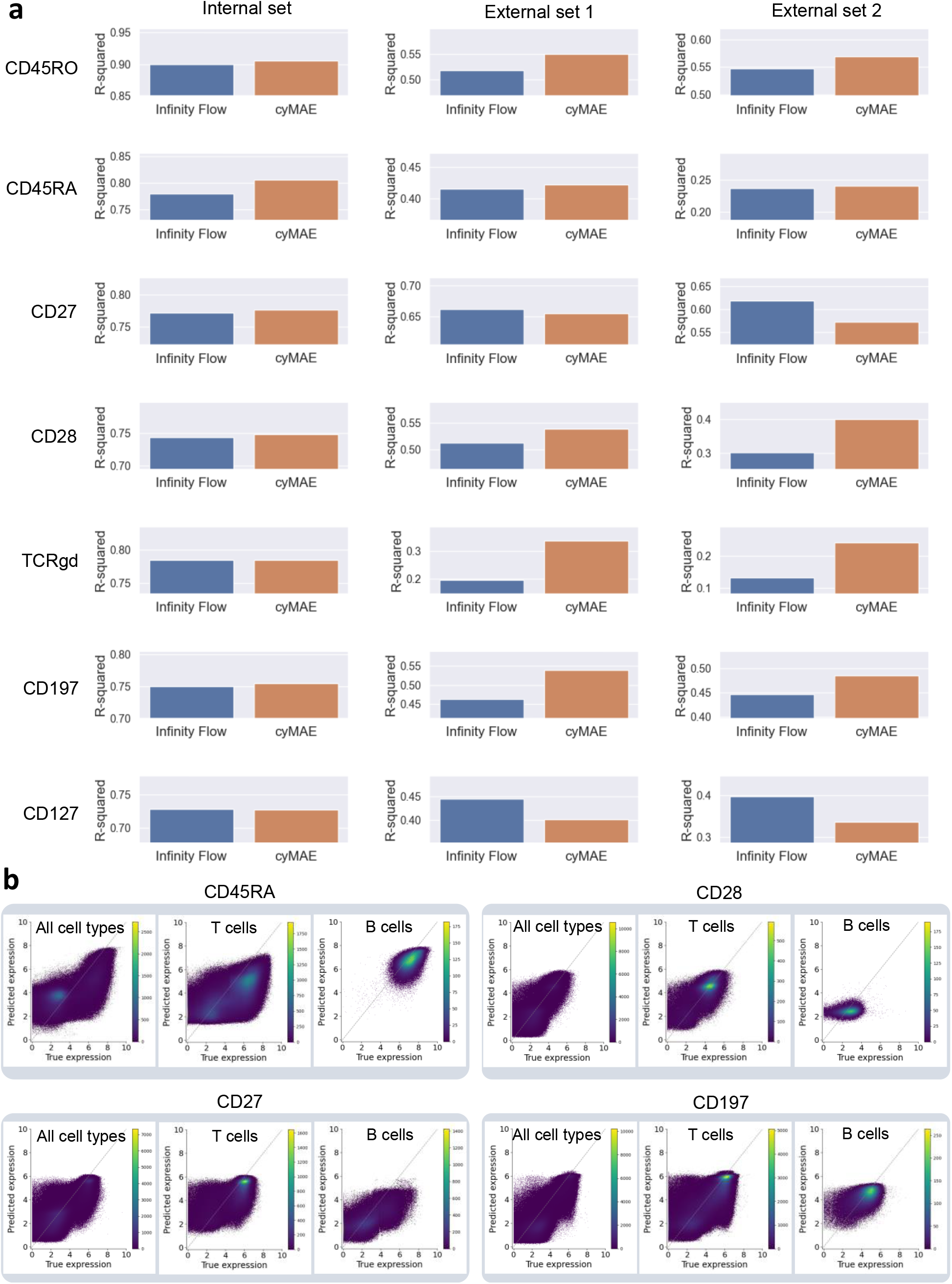
(a) R-squared comparison between Infinity Flow and cyMAE for the imputation task. A total 7 markers were masked and then predicted by the two models. (b) Plots of actual versus predicted expression levels for each marker in the external set (Vaccine dataset). The dashed line represents the ideal relationship, serving as a reference to assess the performance.

### cyMAE is an interpretable immunophenotyper

A key challenge in deploying machine learning models for clinical or biological analyses is their “black box” nature, which means that the rationale for cell type prediction decisions is difficult to interpret. Unlike these models, cyMAE incorporates a multi-head self-attention mechanism within its transformer architecture, enabling interpretable predictions for downstream tasks. The attention scores generated by this model indicate the importance of specific marker information and their interrelations in the context of prediction tasks, with higher attention score indicating greater reliance on a maker’s information relative to other markers. In our analysis, we first measured the attention scores attributed to each feature across different cell types during cell type annotation (**Figure 4a**). Notably, CD45 consistently emerged as the marker with the highest attention score across all cell types, serving as a key discriminator between major immune cell lineages, such as granulocytes and mononuclear cells. Aside from CD45, most markers were highly attended in cell types in which they are highly expressed: for example, CD19 in B cells, CD123 in basophils and pDCs, CD294 in basophils and eosinophils.

**Figure 4.**
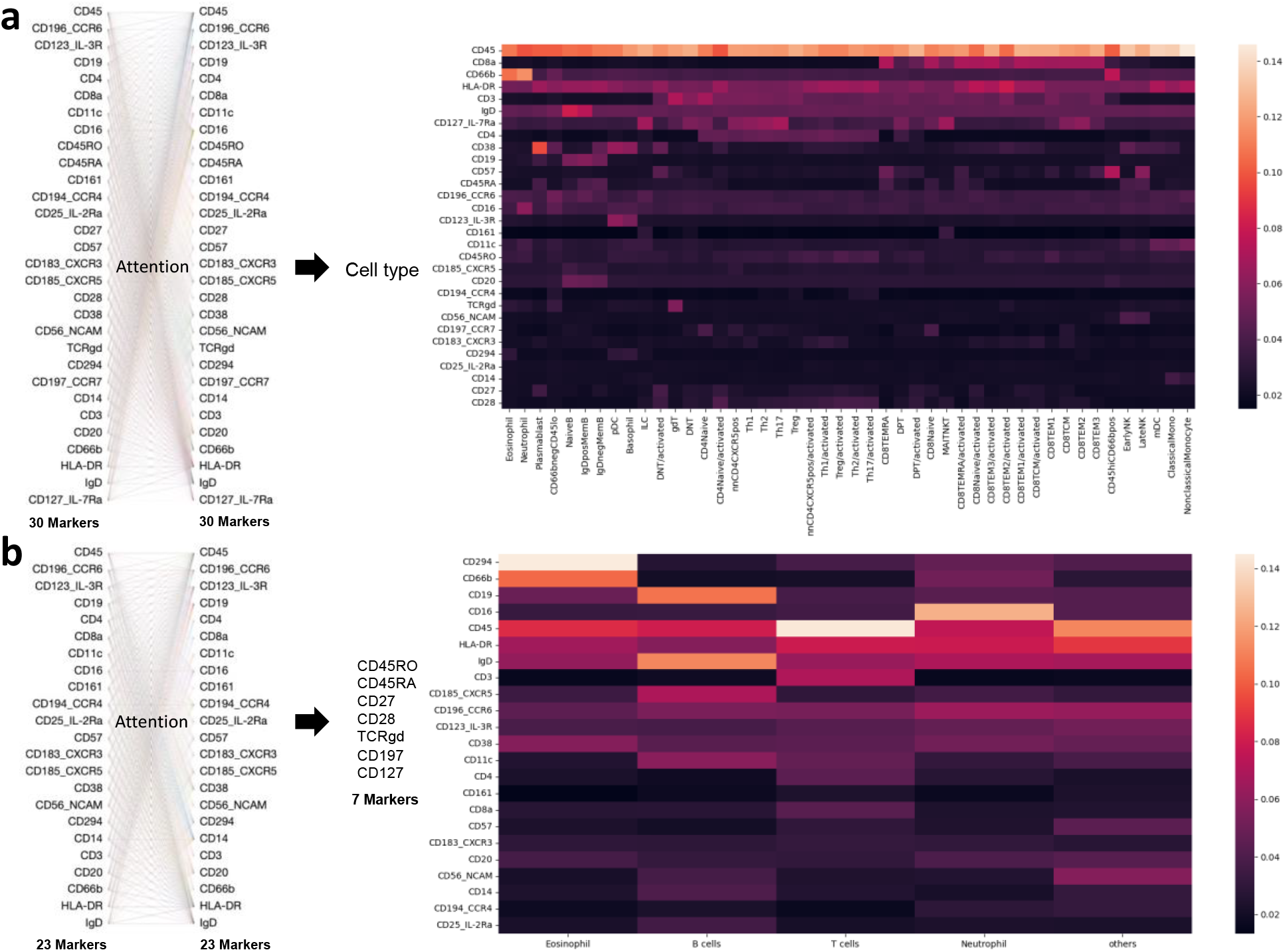
(a) Interpretation in the cell type annotation by the attention scores for the Acute2021 dataset (external set 1). The heatmap shows protein markers with high attention score as bright red for each cell type. (b) Interpretation in the imputation task by the attention score for the Vaccine dataset (external set 1). From 23 markers to impute the other 7 markers, it measures which input features have high attention from the other features during prediction. The heatmap shows the protein markers with high attention score as bright white or red for each cell type. For the left figure in (a) and (b), we used Bertvis^36^ for visualization of attention weights.

Similarly, we assessed the attention score of 23 markers relative to each cell type in the context of predicting the expression of 7 masked markers in the Imputation task (**Figure 4b**). For cell types with constitutive expression or non-expression of masked markers, the model primarily focused on markers indicative of cell type identity (e.g., CD294, CD66b, CD45 for eosinophils; CD16, CD45, and HLA-DR for neutrophils). In the case of T cells, where knowing the cell type was insufficient for predicting expression of the masked proteins, the model attended to the T cell marker protein CD3, but also to CD45 and HLA-DR, both of which were negatively correlated with CD45RA (**Supplementary Figure 12**). The negative correlation between CD45 and CD45RA in T cells was not expected by the authors, but cyMAE found and exploited it to improve imputation performance.

The attention scores not only demonstrated a consistent pattern across external datasets (**Supplementary Figure 9**) but also showed minimal variance between samples (**Supplementary Figure 10**,**11**), underscoring the model’s interpretability and reliability in identifying critical biomarkers for cell type predictions.

### cyMAE improves subject status predictions

In a typical flow cytometry or CyTOF analysis, hundreds of thousands of single cells are generally obtained from an individual sample, aiming to understand cellular-level immunophenotypes, like cell type identification. However, extending these analyses to achieve phenotypic precision at the individual level is also critical. While manual gating is a sophisticated method for extracting subject-level features using expert knowledge, it may overlook complex co-expression patterns indicative of cellular states like activation, senescence, or exhaustion in T cells due to the high-dimensional nature of accurate definintions of these T cell states. Ideally, our aim is to leverage the comprehensive global distribution of cell information to gain deeper biological insights.

A key requirement for this goal is ensuring the method’s predictions remain consistent regardless of the cells’ order in the dataset, a property known as permutation invariance. This property ensures that the method is robust and not reliant on the specific ordering of cells. Additionally, the method should adaptively focus on marker cell types relevant to the study, such as leukemic blast cells in an acute myeloid leukemia (AML) study^34^ or CTLA4+ or PD1+ cells in cancer immunotherapy study^35^. To address these needs, we aggregated cyMAE representations of all cells from each subject into a subject-level representation, exploring several pooling methods to find the most effective one for each task.

We compared the proposed approach with manual gating and FlowSOM on three prediction tasks (**Figure 5a**). Using 5-fold cross-validation with 10 repetitions for each task, we first assessed the ability to distinguish between COVID-19 patients and healthy subjects. Manual gating and FlowSOM showed high accuracy of 0.975 and 0.936 AUROC, respectively, on the test set. In cyMAE, global min pooling performed the best, with an accuracy of 0.987 AUROC (Cohen’s *d* = 0.310 and *d* = 0.706, respectively) (**Supplementary Table 2**). The second challenge was to predict whether a secondary or recall immune response was triggered by SARS-CoV-2 infection or by SARS-CoV-2 vaccination. Manual gating and FlowSOM showed performance of 0.641 and 0.579 AUROC, whereas cyMAE has an AUROC of 0.669 when using global min pooling (*d* = 0.183 and *d* = 0.594, respectively) (Supplementary Table 3). Finally, we tested the ability to distinguish the pre- and post-treatment status of COVID-19 patients. Manual gating and FlowSOM showed 0.796 and 0.869 AUROC, whereas cyMAE showed an accuracy of 0.861 AUROC with global max pooling (*d* = 0.568 and *d* = 0.019, respectively) (**Supplementary Table 4**). The results of the three experiments suggest that cyMAE captures critical information overlooked by manual gating or FlowSOM, thereby enhancing the prediction of subject status across various tasks.

**Figure 5.**
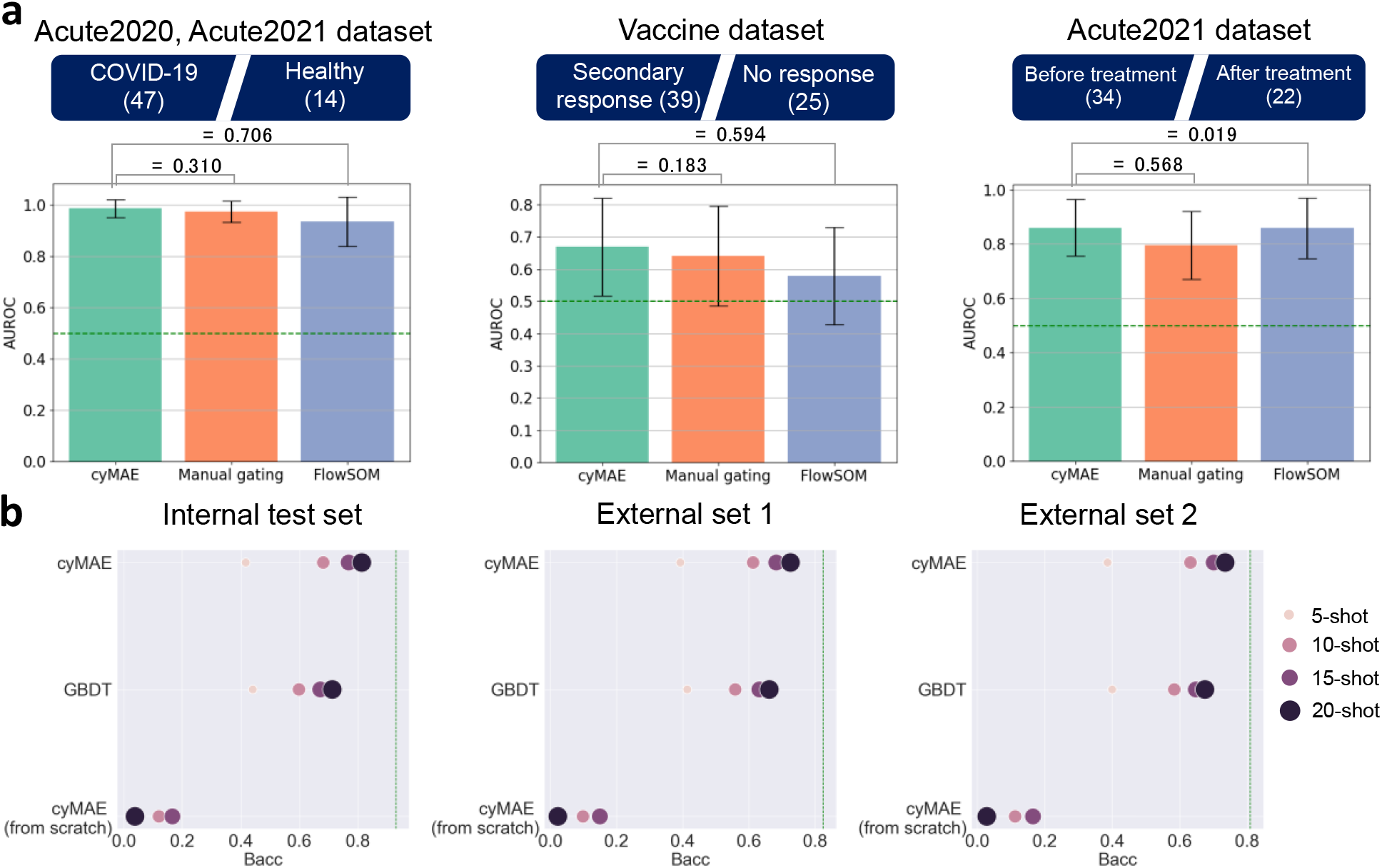
(a) From left to right, COVID-19 patient and healthy people classification using the Acute2020 and Acute2021 dataset, secondary immune response against COVID-19 prediction using the Vaccine dataset, and COVID-19 pre- and post-treatment classification using the Acute2021 dataset. The number in parentheses is the sample size. All the experiments are conducted by 5-fold cross-validation repeating 10 times, and Cohen’s *d* was used to measure the differences between two methods. Green dashed lines stand for performance of a random classifier. (b) The few-shot learning for cell type annotation. Each green dashed line represents the performance of the full fine-tuned cyMAE when used all available training set, reported in the Figure 2b.

### cyMAE is a few-shot learner

Unlike full fine-tuning, few-shot learning trains a model with a limited amount of training data. N-shot uses only N samples for each class in the classification problem. A pre-trained large language model, developed through self-supervised learning, is recognized for its effectiveness as a few-shot learner^11^. Similarly, we evaluated our model, cyMAE, in a few-shot learning context for cell type annotation, conducting experiments with 5-shot, 10-shot, 15-shot and 20-shot settings. Training, validation, testing, and external testing sets are the same as in the previous cell type annotation tasks.

As expected, the performance of cyMAE, when pre-trained, approached that of training with the full training set as the number of N-shots increased (**Figure 5b**). On the other hand, since the cyMAE from scratch (non-pre-trained) has many parameters and no pre-trained information, this method does not learn with small sample size. It is worth noting that GBDT also performed reasonably well, but cyMAE outperformed GBDT based on the pre-trained knowledge. This analysis shows that cyMAE, once pre-trained, can effectively adapt to new tasks even with sparse labeled data, guiding learning in the appropriate direction.

## Discussion

Due to the popularity, ease, and relative affordability of cytometry experiments, there is an abundance of high-dimensional cytometry data compared to other single cell modalities. Although manual gating remains the preferred classification approach, it becomes impractical for the expansive datasets of multi-cohort and/or multi-institutional studies due to its time-consuming and labor-intensive nature. Additionally, some clustering methods, which require loading all the data simultaneously, are not suitable for large-scale datasets due to memory constraints. On the other hand, the cyMAE method uses a mini-batch approach for processing large-scale datasets, where it breaks down the data into small, manageable segment. This approach reduces memory demands and improves training efficiency on large-scale data. The learning time in the training phase scales linearly with the number of samples. Also, cyMAE can quickly and accurately make inferences on new datasets once the model has been trained. In summary, we demonstrate that cyMAE is a scalable solution superior to existing methods for analyzing large-scale data.

To enable a direct comparison of methods, we adopted a paradigm of training models and then using them to make inferences on a new dataset. In contrast, manual gating usually imports historical gates, which are then manually adjusted when necessary for each sample, a time consuming and often error-prone approach. The alternative of simply using clustering approaches to discover sources of variability in each dataset independently can be difficult to scale and does not use a priori information on cell types. The main advantages of the train-inference paradigm are scalability and reproducibility: any investigator can apply the exact same model to any dataset, obtaining results that are easily interpretable within the biologically established framework of immunology. These results show that cyMAE outperforms alternative models within this paradigm.

Directly comparing learning without pre-training (from scratch) and with pre-training, performance improved not only in cell type annotation but also in the few-shot setting (**Supplementary Table S1, Figure 5b**). This model was able to learn stably, while the from-scratch model was unable to learn effectively with little training data. While not a dramatic performance improvement, in the other experiments using the pre-trained cell embeddings, it was encouraging to see that the pre-trained embedding was good at learning antibody co-localization patterns, imputing unavailable protein expressions, and contributes meaningful performance gains for the subject-level predictions.

In this study, we pre-trained our model using only one of the three available cohorts to evaluate the performance on several downstream tasks using all three cohorts. Future research will expand this approach by pre-training on a broader array of data from multiple studies, including more diverse subject phenotypes. This expansion is expected to enhance the model’s power and robustness, enabling it to more effectively distinguish between biological variations and gain a deeper understanding of protein functions and protein expression patterns. This, in turn, will lead to more accurate predictions in various downstream tasks.

This study has several limitations. First, while these models are trained on only CyTOF data, its application to flow cytometry data might not be recommended due to inherent technical differences. Specifically, the methodologies used in flow and mass cytometry yield disparate patterns of protein expression. Yet, a model like cyMAE, if pre-trained on flow cytometry data from scratch, could indeed become a viable approach for flow cytometry datasets. Second, we assume that cell type information from manual gating is the ground truth. However, this may not be the case in practice. Even the most skilled experts are prone to subjectivity and bias, which might lead to a bias toward “expected” results. This claim can be reinforced by our experimental results of the subject status prediction, where the pre-trained cyMAE showed higher predictive power than the manual gated features in some tasks (**Figure 5a**). This observation raises the possibility that there may be information that manual gating misses. Finally, the size and type of cytometry panels used in practice vary widely depending on the research purpose. However, our model was only trained to work on data with fixed markers. For more meaningful research, it should work robustly for different panels in future studies, for examples, to accommodate data where only a subset of the markers has been measured. Despite these limitations, this study demonstrates the high potential of pre-training in single-cell cytometry, both because an approach like ours has not been applied to cytometry data analysis before and because it shows advantages over previous methods.

Here, we introduced cyMAE, a masked autoencoder model which builds latent embeddings of single-cell cytometry data and uses them to achieve good performance across a range of cell-level and subject-level tasks. Especially, the fine-tuned cyMAE is as accurate as manual gating, with the labor-free advantages of automated analysis. To the best of our knowledge, cyMAE is the first such model which specializes on cytometry data. Our results are a proof of concept for applying a combination of unsupervised and supervised analysis in the training-inference paradigm to multiple cytometry datasets that use the same panel. This approach promises scalability across thousands of samples from multiple studies, providing robust and interpretable results while minimizing manual analysis.

## Methods

### Human Subjects

All subjects consented and enrolled with approval of the University of Pennsylvania Institutional Review Board (Vaccine IRB no. 844642; Acute2020 IRB no. 808542; Acute2021 IRB no. 843758). All participants or their surrogates provided informed consent in accordance with protocols approved by the regional ethical research boards and the Declaration of Helsinki.

For the Vaccine dataset, 43 individuals were enrolled for longitudinal monitoring of response to SARS-CoV-2 mRNA vaccine beginning in December 2020 through March 2021. All subjects received either Pfizer (BNT162b2) or Moderna (mRNA-1273) mRNA vaccines. Samples were collected at six time points: baseline, ∼2 weeks after primary immunization, day of secondary immunization, ∼1 week after secondary immunization, ∼3 months after primary immunization, and ∼6 months after primary immunization. Participants were self-reported healthy without ongoing chronic health conditions. In the Vaccine dataset, the definition of secondary immune response was defined as follows. We labeled a secondary immune response as “Yes” if it occurred after a healthy person received two vaccines, or after a person with COVID-19 received one vaccine, or after a person with COVID-19 received two vaccines. If a healthy person received a single vaccine, we labeled it “No”.

For the Acute2020 dataset, patients were consented and enrolled within 3 days of admission to the Hospital of the University of Pennsylvania with a positive SARS-CoV-2 PCR test, regardless of the oxygen support needed. Clinical data were abstracted from the electronic medical record into standardized case report forms. All subjects in this dataset were consented and enrolled between March and December 2020 at the University of Pennsylvania. Subjects in the Acute2021 dataset were enrolled in the I-SPY Covid-19 Trial^28^. Hospitalized participants at 5 trial sites (Penn, University of Alabama Birmingham, University of California San Francisco, University of Colorado, and Wake Forest University Atrium Health) with confirmed SARS-CoV-2 PCR or antigen testing and requiring greater than 6 liter per minute oxygen flow (including high flow nasal oxygen, high flow face mask oxygen, non-invasive ventilation, or invasive mechanical ventilation consistent with World Health Organization ordinal scale ≥ 5) for fewer than 72 hours were enrolled in this trial. Patients or their legally authorized representatives consented to be randomized to receive a backbone treatment (remdesivir and dexamethasone) alone versus backbone with one of 12 investigational treatments. Details of the trial inclusion and exclusion criteria, and the non-backbone treatment arms have been published at https://clinicaltrials.gov/study/NCT04488081. Whole blood was collected at time of admission and 7 days later. Samples from subjects enrolled at the University of Pennsylvania were processed on the day of collection. Samples from subjects enrolled at the University of Alabama at Birmingham, University of Colorado, University of California at San Francisco, and Wake Forest University were shipped to the University of Pennsylvania and processed the day of arrival.

### Mass Cytometry

For all samples, 270μL of whole blood were stained using the MaxPar Direct Immune Profiling Assay (Standard BioTools, Inc, South San Francisco, CA)^37^. For the Acute2020 dataset, samples were stained in accordance with manufacturer protocols. Briefly, whole blood was added to a 5mL tube containing a pellet of lyophilized antibodies. Blood was incubated at room temperature for 30 minutes and then lysed with Cal-Lyse lysing solution Standard BioTools, Inc, South San Francisco, CA). Cells were washed, followed by fixation with 1.6% PFA. Cells sat at 4°C over night prior to staining with Cell-ID Intercalator-Ir. These samples are referred to as “fresh” because they did not undergo cryopreservation and thawing. Vaccine and Acute2021 data sets underwent a similar workflow as described above. However, after incubating for 30 minutes in the tube of lyophilized antibodies, stained whole blood was fixed with PROT1 buffer (Smart Tube Inc, Las Vegas, NV) and cryopreserved. Lyse, wash, and intercalator staining were performed as above after thaw. Stained samples were collected on a CyTOF 2 instrument with EQ4 beads (four element calibration beads, Standard BioTools, Inc).

After data acquisition, .fcs files were gated to remove beads, debris, doublets, and dead cells using the OMIQ platform (Boston, MA); representative gates are shown in Supplementary Figure 13. After gating, DNA intercalator, viability, Gaussian and bead channels were dropped, and the remaining protein expression channels were transformed using inverse hyperbolic sine with a cofactor of 5.

### Model details

#### Transformer block

The transformer block consists of alternating layers of multihead self-attention (MSA) and multilayer perceptron (MLP) blocks (Equation 1,2). Layer norm (LN)^38^ is applied before every block, and Drop path (DP)^39^ is applied after every block. The MLP contains two linear layers with GELU activation function.

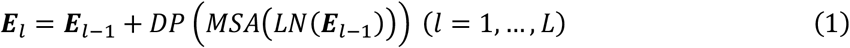

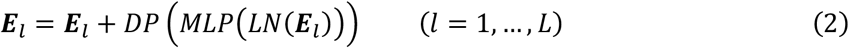

where ***E***_*l*−1_ denotes output embeddings of the (*l* − 1)-th layer and input embeddings of the *l*-th layer at the same time.

#### Multi-head self-attention

In the multi-head self-attention (MSA) layer, we compute query, key, and value matrix (***Q, K, V***) from the input embeddings (***E***) for each head (Equation 3) and compute *h* heads by weighted sum of all values by attention weight for each head, where attention weight is calculated by the pairwise similarity between two elements of the input and their respective query and key representations (Equation 4). Finally, *h* heads are concatenated, and the output is linearly projected (Equation 5)

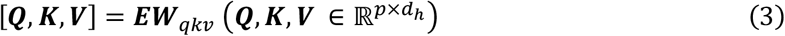

where ***E*** ∈ ℝ^*p*×*d*^ is input embeddings 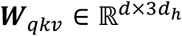 is learnable weight matrix, and *d*_*h*_ is set to *d*/*h*.

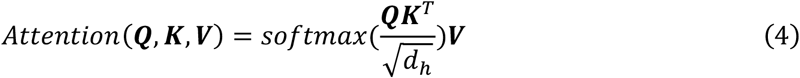

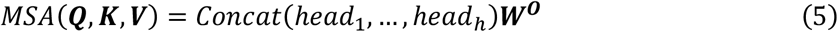

where *head*_*i*_ = *Attention*(*Q*_*i*_, *K*_*i*_, *V*_*i*_)(*i* = 1, …, *h*), and ***W***^***O***^ ∈ ℝ^*d*×*d*^ is linear weight matrix.

#### Cytometry Masked Autoencoder

The whole structure consists of an encoder and a decoder, which are used in the pre-training step. The encoder is only then used with a single linear layer in the downstream supervised fine-tuning. The encoder (*f*_*e*_) consists of 12 layers of transformer blocks, each with 12 heads and 768 hidden dimensions, totaling 85 million parameters. In contrast, the decoder (*f*_*d*_) is smaller than the encoder. It consists of 4 layers of transformer blocks with 6 heads and 384 hidden dimensions for a total of 7 million parameters. The dimension size of latent cell or subject representations for the downstream tasks is 768. This setting was proposed in the Masked autoencoder.

### Training details

#### Masked Cytometry Modeling (MCM)

cyMAE learns to maximize

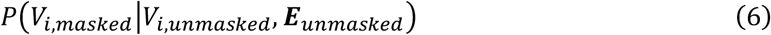

for cell *i*, where *V*_*i*,***mask****ed*_ ∈ ℝ ^(*r*·*p*)×1^ denotes masked protein expressions of cell *i*, and ***E***_*masked*_ ∈ ℝ ^(*r*·*p*)×(*d*−1)^ denotes masked protein embeddings after masking. *r* is a masking ratio, *p* is the number of proteins in the data, and *d* is a hidden dimension size. Likewise, *V*_*i*,***un****masked*_ ∈ ℝ ^(1−*r*)·*p*×1^ denotes unmasked protein expressions of cell *i*, and ***E***_*unmasked*_ ∈ ℝ ^(1−*r*)·*p*×(*d*−1)^ denotes unmasked protein embeddings.

The encoder (*f*_*e*_) generates a latent representation of the cell. The unmasked latent representation of cell *i* is defined as ***H***_*i*,***un****masked*_ ∈ ℝ ^(1−*r*)·*p*×*d*^ as the following,

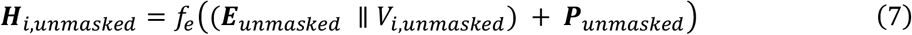

where ***P***_*unmasked*_ ∈ ℝ ^(1−*r*)·*p*×*d*^ is sine-cosine positional embeddings for masked proteins. The idea of the concatenation (∥) of protein embeddings with expression values was inspired from MET^40^.

The decoder (*f*_*d*_) reconstructs the masked values as followings,

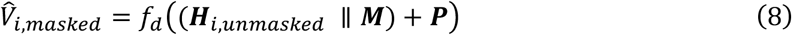

Let *M* denote a learnable mask token embedding represented as a row vector *M* ∈ ℝ^1×*d*^. We construct a matrix ***M*** by stacking this vector *r* · *p* times, such that the resulting matrix ***M*** has dimensions (*r* · *p* × *d*). ***P*** ∈ ℝ^*p*×*d*^ is sine-cosine positional embeddings. To calculate the reconstruction loss, we use mean square error (MSE) loss for all cells,

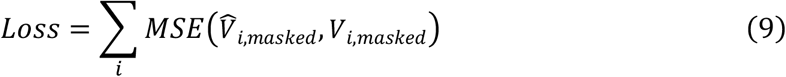

#### Why positional embedding is necessary

It might seem that positional embedding is not necessary because the input is a tabular data. However, the position serves as an index to indicate which protein’s expression value should be reconstructed by the decoder during *MCM*. For example, 2nd, 3rd, and 7th proteins of 10 proteins are masked, positional embedding provides information to reconstruct the expression of the 2nd, 3rd, and 7th proteins. Therefore, when using cyMAE, users make sure to match the order of the proteins.

#### Cell representation

After pre-training, the cell representation (***C***_*i*_) of cell *i* is obtained as follows.

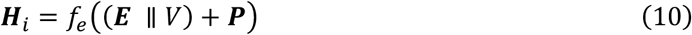

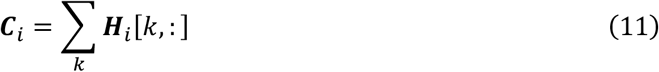

Then, this cell representation is used as input of a linear layer for cell-level downstream tasks.

#### Subject representation

The subject representation is obtained by multiple global pooling methods.

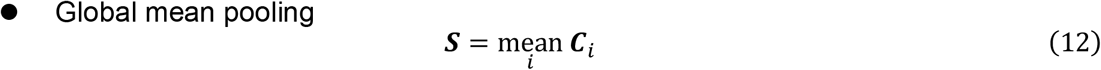

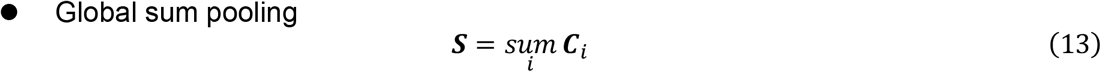

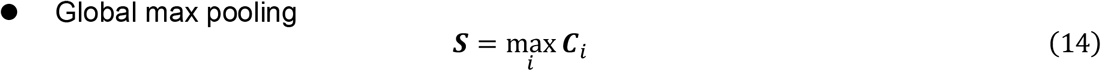

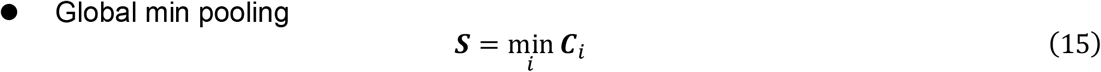

Then, this subject representation is used as input of a linear layer for subject-level supervised downstream tasks.

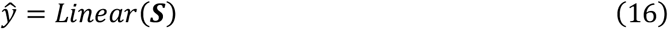

#### Supervised learning in downstream tasks

Cross entropy loss is employed for classification tasks and Mean squared error (MSE) loss is employed for regression tasks.

In the cell type annotation task,

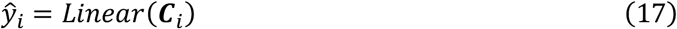

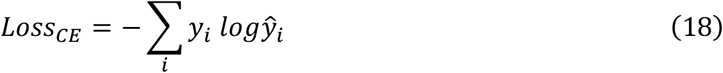

where *y*_*i*_ and ŷ_*i*_ indicate the ground truth cell type and the predicted probability for cell type of cell *i*, respectively.

In the imputation task,

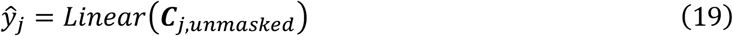

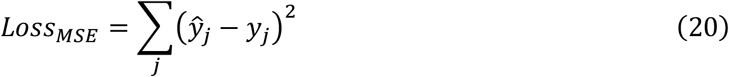

where *y*_*j*_ and ŷ_*j*_ denote the ground truth expression value and the predicted value of masked protein *j*, respectively.

In the subject-level prediction tasks,

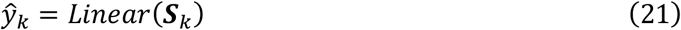

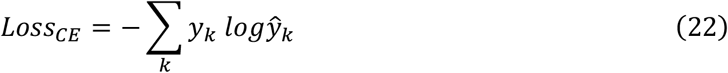

where *y*_*k*_ and ŷ_*k*_ are the ground truth label and predicted probability for label of subject *k*, respectively.

#### Impact of Masking ratio during pre-training

To test if masking ratio affects cyMAE training, we trained three different versions of the model with masking ratios of 0.25, 0.5, and 0.75. The result was there was no significant difference in performance on the cell type annotation tasks (**Supplementary Table 5**). Therefore, all the cyMAE experiments were performed with a 0.25 masking ratio.

#### Training setting

The configuration includes a batch size of 768, drop path regularization of 0.1, AdamW optimizer with momentum of 0.9 and weight decay of 0.05, learning rate of 0.0005 with a cosine scheduler, and label smoothing during fine-tuning.

#### Computational cost in training and inference

The pre-training required 10 days with four of GeForce RTX 2080 Ti Rev. A to process 6.5M cells through 200 epochs. Fine-tuning the model for cell type annotation took 13 days on a single GeForce RTX 2080 Ti Rev. A GPUs to process 29.4 million cells through 100 epochs, with early stopping implemented. For inference, the runtime was 1.2 hours for 7.3M cells under the Vaccine dataset and 2.1 hours for 18.4M cells under the Acute2020 dataset and Acute2021 dataset, both on a single GPU.

### Benchmarking models

#### Manual gating

Each sample from all datasets was manually gated using the OMIQ platform to obtain the 46 terminal populations used as ground truth labels. Representative gates from our strategy are shown in **Supplementary Figure 3**.

#### Static gating

For each gate in our hierarchy, we aggregated the candidate gate positions from all training samples in the Vaccine dataset into one consensus gate. By definition, a point is in the consensus gate if it falls into at least 30% of all the candidate gates (Supplementary Figure 4). We then created a consensus hierarchy out of all consensus gates and applied it statically to all test samples.

#### FlowSOM clustering

The same 60% of the Vaccine data samples were used to train an unsupervised FlowSOM clustering model. Version 2.6.0 of the FlowSOM R package was used with default parameters, except for the total number of metaclusters, which we set to 46 to match the number of ground truth labels. As an unsupervised clustering algorithm, FlowSOM does not have an inference mode. We performed inference on testing datasets (20% of the Vaccine dataset as an internal test set, and the two external test sets) by assigning each datapoint to the nearest SOM node from the trained model, and preserving the assignment of nodes to metaclusters from the training phase. Evaluation of accuracy and balanced accuracy required the extra information of a bipartite matching between the 46 FlowSOM clusters and the 46 ground truth labels. Following Weber, L. M. *et al*.^41^, we obtained the matching using the Hungarian algorithm, implemented in the function *solve_LSAP* of the R package clue.

#### Gradient Boosting Decision Tree (GBDT)

We used XGBoost^32^ python package for GBDT. We ran XGBoost regressor or classifier with 100 estimators and 0.03 learning rate and set early stopping based on the performance change for the validation set.

#### Fully connected Deep Neural Network (DNN)

DGCyTOF (Cheng, L. *et al*.^30^) and DeepCyTOF (Li, H *et al*.^*31*^) proposed a fully-connected neural network for cytometry data. Both were designed for a cell representation and cell-level prediction tasks, so we use this architecture as a comparison model.

#### Convolutional Neural Network (CNN)

Hu. Z *et al*.^8^ proposed a model using convolutional neural network for cytomegalovirus (CMV) classification. The original model was designed for subject-level tasks, but since it uses a CNN structure to draw cell representations and pool them, we modified to the same architecture without the pooling layer as a comparison model.

### Metrics

#### Balanced accuracy (Bacc)

For a multi-class imbalanced dataset, we used Balanced accuracy (Bacc) instead of Accuracy. Balanced accuracy is defined as a macro-average of recall scores per class in a multi-class classification.

A Recall score is defined as:

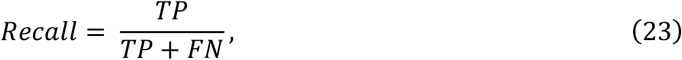

where TP is true positive, and FN is false negative.

#### R-squared

In a regression task, if ŷ_*i*_ is the predicted value of the *i*-th sample and *y*_*i*_ is the corresponding true value for total *n* samples, the R-squared is defined as:

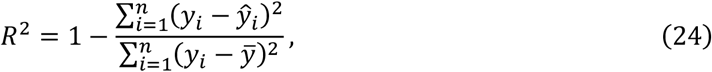

where 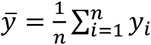.

#### AUROC

A receiver operating characteristic (ROC) curve is widely used for evaluating prediction models. It plots True Positive Rate (TPR) against False Positive Rate (FPR).

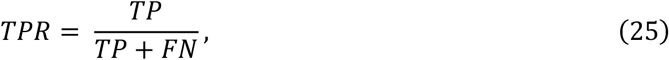

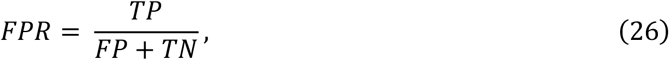

Where TP, FP, TN, and FN are the number of true positives, false positives, true negatives, and false negatives respectively. AUROC stands for the area under the ROC curve.

#### Cohen’s *d*

Cohen’s *d* is defined as the difference between two means divided by a standard deviation for the data,

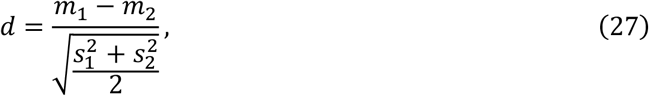

where *m*_1_ and *s*_1_ are mean and standard deviation of method 1 and, *m*_2_ and *s*_2_ are mean and standard deviation of method 2.

### Protein embeddings

After pre-training through *MCM*, the trained ***E*** ∈ ℝ^*p*×(*d*−1)^ in cyMAE represents protein embeddings for *p* proteins. It is expected that they have protein information about the heterogenous, complex, and dynamic immune cells without any prior information, only through learning on the data itself. This was confirmed by the PCA 2-dimensional plot in **Figure 2a**.

### Imputation

We masked CD45RO, CD45RA, CD27, CD28, TCRgd, CD197, and CD127 expressions and used the remaining markers to predict the expression of these seven marker expressions. Infinity Flow used GBDT as the imputer. Similarly, the unmasked cell representations were first extracted from the pre-trained cyMAE and used as input to GBDT to train and then evaluated on the external test sets (not end-to-end).

### Attention score

From Equation 4, we first obtain the output of *softmax* function for the interpretation for cell *i* (Equation 28) and calculate the attention score *W*_*i*_ by averaging over the query axis (Equation 29).

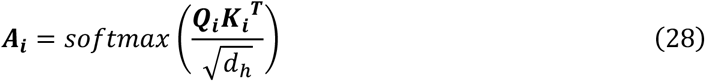

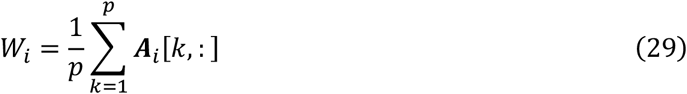

In our experiments, we sampled 2% of all cells for each dataset. To calculate the attention score, we only used the information from the first layer, because the first layer is the most influential in determining which inputs to give attention to, and there was no significant difference in attention between inputs after the second layer.

### Few-shot learning setting in the cell type annotation

For *N*-shots, we trained using only the first *s* samples per class in the training and validation sets and then evaluated on the entire test set. We compared performance for 5, 10, 15, and 20 shots.

### Software

This project would not have been possible without numerous open-source Python packages including torch, torchvision, timm, deepspeed, einops, jupyter, matplotlib, numpy, pandas, scikit-learn, seaborn, FlowCytometryTools, scipy, etc. Specific versions for each package can be found at https://github.com/JaesikKim/cyMAE/blob/master/requirements.txt.

## Data availability

The data presented within this study is available upon formal request to the corresponding author.

## Code availability

All software for dataset construction, model training, deployment, and analysis is available on our Github page: https://github.com/JaesikKim/cyMAE.

## Figure Preparation

All plots were generated via Python scripts using the following open-source visualization packages: seaborn 0.12.1. Microsoft PowerPoint was used for the final annotation and assembly of panels. BioRender was used for generating Figure S2. Mathematical equations were prepared with LaTeX.

## Acknowledgements

We thank Takuya Ohtani and the Penn CyTOF Core at the University of Pennsylvania for data acquisition.

## Author contributions

J.K., M.I., A.R.G., and D.K. conceived the study. J.K. and M.I. conducted analysis with the help of M.L., M.L.M, Y.N., M.M.P., J.W., S.A.A., P.O., S.-H.J., and J.W., and C.C., M.L., and M.L.M. provided critical comments for model development. A.P. conducted sample preparation, and D.T.N. annotated the data by manual gating. Following researchers recruited subjects, collected clinical data, obtained and shipped biosamples for the I-SPY clinical trial: C.A.G.I., A.P.T., M.E., T.G.D., N.S.M., J.P.R., and N.J.M. from Perelman School of Medicine, University of Pennsylvania. C.S.C., K.D.L., M.A.M., and L.B.S. from University of California, San Francisco School of Medicine. E.L.B. and J.M. from University of Colorado School of Medicine. S.G. and D.W.R. from University of Alabama at Birmingham. D.C.F., K.W.G., K.W.T., and H.B. from Wake Forest School of Medicine. J.K., M.I., A.R.G., and Y.N. provided graphical design of figures. J.K. and M.I. wrote the manuscript, and D.M., V.T., A.R.G., D.K., and E.J.W. provided critical comments and valuable edits. A.R.G., D.K., and E.J.W. supervised the study.

## Reference

1 Maecker, H. T., McCoy, J. P. & Nussenblatt, R. Standardizing immunophenotyping for the Human Immunology Project. Nat Rev Immunol 12, 191–200 (2012). 10.1038/nri3158

2 Mair, F. et al. The end of gating? An introduction to automated analysis of high dimensional cytometry data. Eur J Immunol 46, 34–43 (2016). 10.1002/eji.201545774

3 Olsen, L. R., Leipold, M. D., Pedersen, C. B. & Maecker, H. T. The anatomy of single cell mass cytometry data. Cytometry A 95, 156–172 (2019). 10.1002/cyto.a.23621

4 Van Gassen, S. et al. FlowSOM: Using self-organizing maps for visualization and interpretation of cytometry data. Cytometry A 87, 636–645 (2015). 10.1002/cyto.a.22625

5 Levine, J. H. et al. Data-Driven Phenotypic Dissection of AML Reveals Progenitor-like Cells that Correlate with Prognosis. Cell 162, 184–197 (2015). 10.1016/j.cell.2015.05.047

6 Spitzer, M. H. et al. IMMUNOLOGY. An interactive reference framework for modeling a dynamic immune system. Science 349, 1259425 (2015). 10.1126/science.1259425

7 Samusik, N., Good, Z., Spitzer, M. H., Davis, K. L. & Nolan, G. P. Automated mapping of phenotype space with single-cell data. Nat Methods 13, 493–496 (2016). 10.1038/nmeth.3863

8 Hu, Z., Tang, A., Singh, J., Bhattacharya, S. & Butte, A. J. A robust and interpretable end-to-end deep learning model for cytometry data. Proc Natl Acad Sci U S A 117, 21373–21380 (2020). 10.1073/pnas.2003026117

9 Vaswani, A. et al. Attention is All you Need. Advances in Neural Information Processing Systems 30 (2017).

10 Devlin, J., Chang, M.-W., Lee, K. & Toutanova, K. BERT: Pre-training of Deep Bidirectional Transformers for Language Understanding. Proceedings of the 2019 Conference of the North American Chapter of the Association for Computational Linguistics: Human Language Technologies 1 (2019).

11 Brown, T. et al. Language Models are Few-Shot Learners. Advances in Neural Information Processing Systems 33, 1877--1901 (2020).

12 Dosovitskiy, A. et al. An Image is Worth 16×16 Words: Transformers for Image Recognition at Scale. International Conference on Learning Representations (2021).

13 Bao, H., Dong, L., Piao, S. & Wei, F. BEiT: BERT Pre-Training of Image Transformers. International Conference on Learning Representations (2022).

14 He, K. et al. Masked Autoencoders Are Scalable Vision Learners. Proceedings of the IEEE/CVF Conference on Computer Vision and Pattern Recognition (CVPR), 16000–16009 (2022).

15 Madani, A. et al. Large language models generate functional protein sequences across diverse families. Nat Biotechnol (2023). 10.1038/s41587-022-01618-2

16 Elnaggar, A. et al. ProtTrans: Toward Understanding the Language of Life Through Self-Supervised Learning. Ieee T Pattern Anal 44, 7112–7127 (2022). 10.1109/Tpami.2021.3095381

17 Lin, Z. et al. Evolutionary-scale prediction of atomic-level protein structure with a language model. Science 379, 1123–1130 (2023). 10.1126/science.ade2574

18 Hie, B. L. et al. Efficient evolution of human antibodies from general protein language models. Nat Biotechnol (2023). 10.1038/s41587-023-01763-2

19 Shanehsazzadeh, A. et al. Unlocking de novo antibody design with generative artificial intelligence. bioRxiv, 2023.2001.2008.523187 (2023). 10.1101/2023.01.08.523187

20 Eguchi, R. R. et al. Deep Generative Design of Epitope-Specific Binding Proteins by Latent Conformation Optimization. bioRxiv, 2022.2012.2022.521698 (2022). 10.1101/2022.12.22.521698

21 Cui, H., Wang, C., Maan, H. & Wang, B. scGPT: Towards Building a Foundation Model for Single-Cell Multi-omics Using Generative AI. bioRxiv, 2023.2004.2030.538439 (2023). 10.1101/2023.04.30.538439

22 Gong, J. et al. xTrimoGene: An Efficient and Scalable Representation Learner for Single-Cell RNA-Seq Data. bioRxiv, 2023.2003.2024.534055 (2023). 10.1101/2023.03.24.534055

23 Yang, F. et al. scBERT as a large-scale pretrained deep language model for cell type annotation of single-cell RNA-seq data. Nat Mach Intell 4, 852-+ (2022). 10.1038/s42256-022-00534-z

24 Chen, J. et al. Transformer for one stop interpretable cell type annotation. Nat Commun 14, 223 (2023). 10.1038/s41467-023-35923-4

25 Shen, H. et al. Generative pretraining from large-scale transcriptomes for single-cell deciphering. iScience 26, 106536 (2023). 10.1016/j.isci.2023.106536

26 Minsheng Hao, J. G., Xin Zeng, Chiming Liu, Yucheng Guo, Xingyi Cheng, Taifeng Wang, Jianzhu Ma, L. Song, View ORCID ProfileXuegong Zhang. Large Scale Foundation Model on Single-cell Transcriptomics. bioRxiv (2023). 10.1101/2023.05.29.542705

27 Theodoris, C. V. et al. Transfer learning enables predictions in network biology. Nature 618, 616–624 (2023). 10.1038/s41586-023-06139-9

28 Consortium, I. S. C. Report of the first seven agents in the I-SPY COVID trial: a phase 2, open label, adaptive platform randomised controlled trial. EClinicalMedicine 58, 101889 (2023). 10.1016/j.eclinm.2023.101889

29 Mathew, D. et al. Deep immune profiling of COVID-19 patients reveals distinct immunotypes with therapeutic implications. Science 369 (2020). 10.1126/science.abc8511

30 Cheng, L., Karkhanis, P., Gokbag, B., Liu, Y. & Li, L. DGCyTOF: Deep learning with graphic cluster visualization to predict cell types of single cell mass cytometry data. PLoS Comput Biol 18, e1008885 (2022). 10.1371/journal.pcbi.1008885

31 Li, H. et al. Gating mass cytometry data by deep learning. Bioinformatics 33, 3423–3430 (2017). 10.1093/bioinformatics/btx448

32 Ester, M., Kriegel, H. P. & Xu, X. XGBoost: A scalable tree boosting system. In Proceedings of the 22Nd ACM SIGKDD International Conference on Knowledge Discovery and Data Mining (vol, pg 785, 2016). Geogr Anal (2022). 10.1111/gean.12315

33 Becht, E. et al. High-throughput single-cell quantification of hundreds of proteins using conventional flow cytometry and machine learning. Sci Adv 7, eabg0505 (2021). 10.1126/sciadv.abg0505

34 Lowenberg, B., Downing, J. R. & Burnett, A. Acute myeloid leukemia. N Engl J Med 341, 1051–1062 (1999). 10.1056/NEJM199909303411407

35 Pardoll, D. M. The blockade of immune checkpoints in cancer immunotherapy. Nat Rev Cancer 12, 252–264 (2012). 10.1038/nrc3239

36 Vig, J. A Multiscale Visualization of Attention in the Transformer Model. Proceedings of the 57th Annual Meeting of the Association for Computational Linguistics: System Demonstrations, 37—42 (2019).

37 Geanon, D. et al. A streamlined whole blood CyTOF workflow defines a circulating immune cell signature of COVID-19. Cytometry A 99, 446–461 (2021). 10.1002/cyto.a.24317

38 Ba, J. L., Kiros, J. R. & Hinton, G. E. Layer Normalization. arXiv (2016).

39 Larsson, G., Maire, M. & Shakhnarovich, G. FractalNet: Ultra-Deep Neural Networks without Residuals. International Conference on Learning Representations (2017).

40 Majmundar, K. A., Goyal, S., Netrapalli, P. & Jain, P. MET: Masked Encoding for Tabular Data. NeurIPS 2022 First Table Representation Workshop (2022).

41 Weber, L. M. & Robinson, M. D. Comparison of clustering methods for high-dimensional single-cell flow and mass cytometry data. Cytometry A 89, 1084–1096 (2016). 10.1002/cyto.a.23030

