## Supplementary materials for "Cytometry Masked Autoencoder: An Accurate and Interpretable Automated Immunophenotyper"

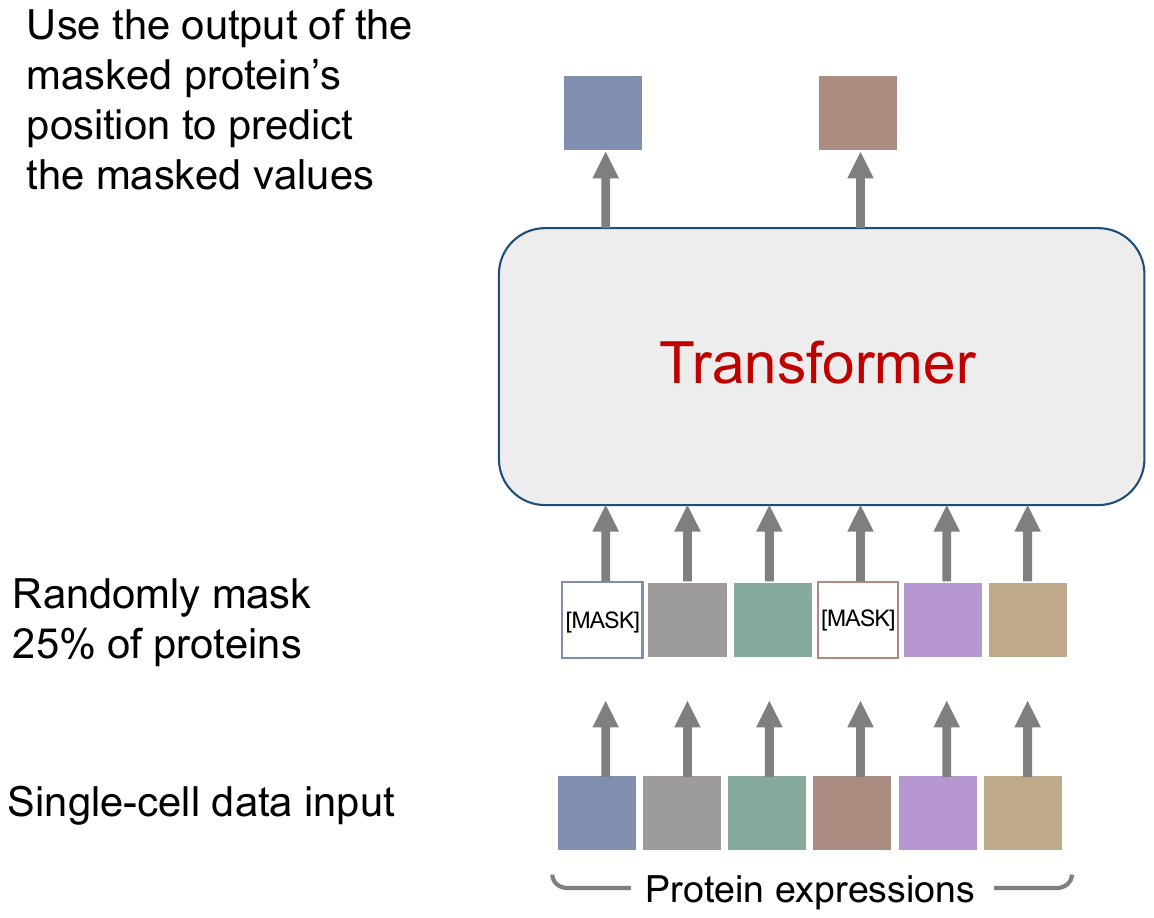

Figure S1. Overview of Masked Cytometry Modelling (*MCM*)

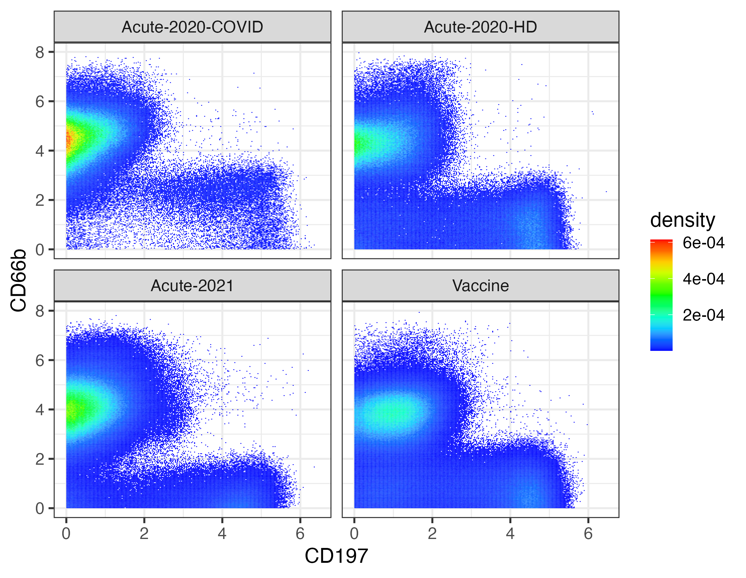

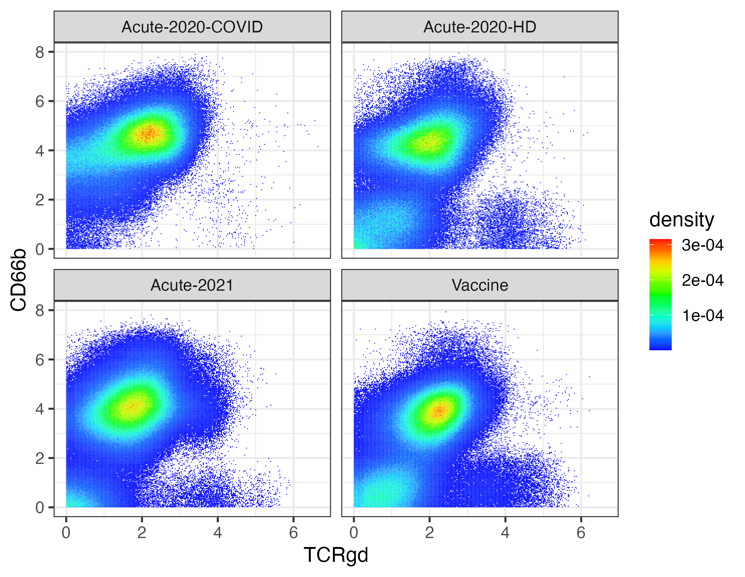

Figure S2. Density plots showing bivariate CD197/CD66b distribution (left) and bivariate TCRgd/CD66b distribution (right) by dataset. Neutrophils and T cells from all samples in a dataset were pooled together; neutrophils are CD66b high and T cells are CD66b low. For the Acute-2020 dataset, the COVID subjects and healthy donors (HD) were plotted separately.

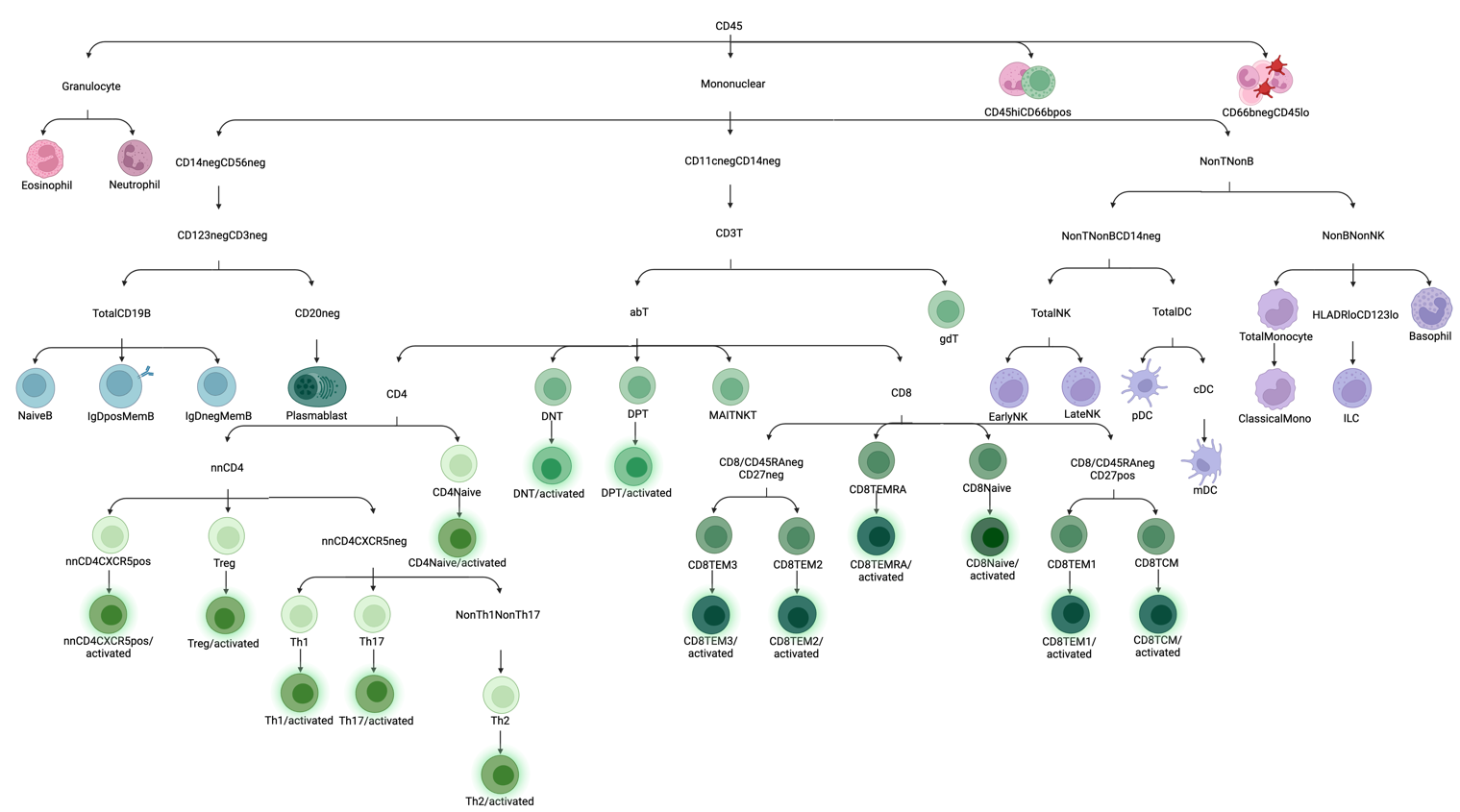

Figure S3. Standard gating strategies for 46 cell types

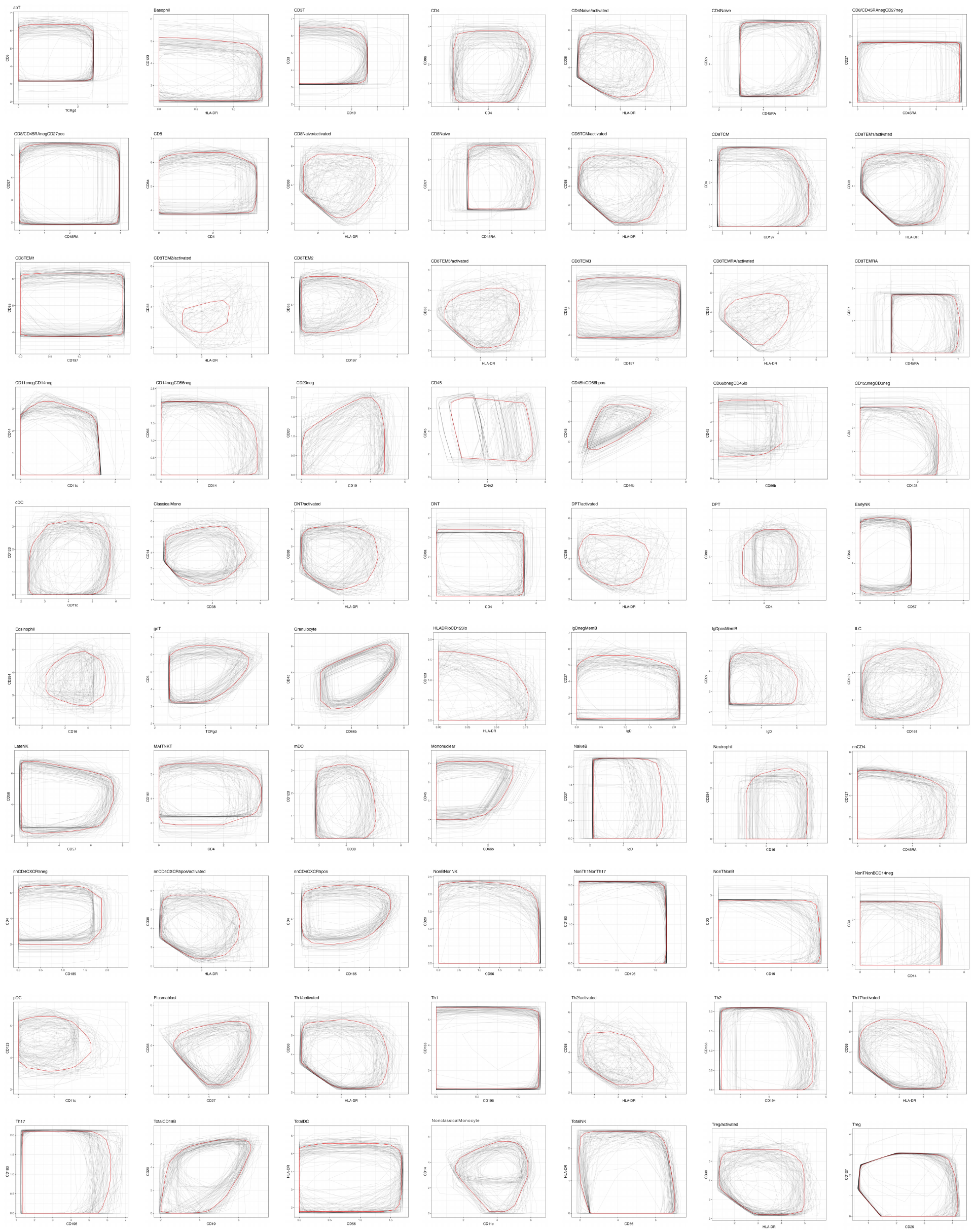

Figure S4. Consensus gates of statical approach in manual gating

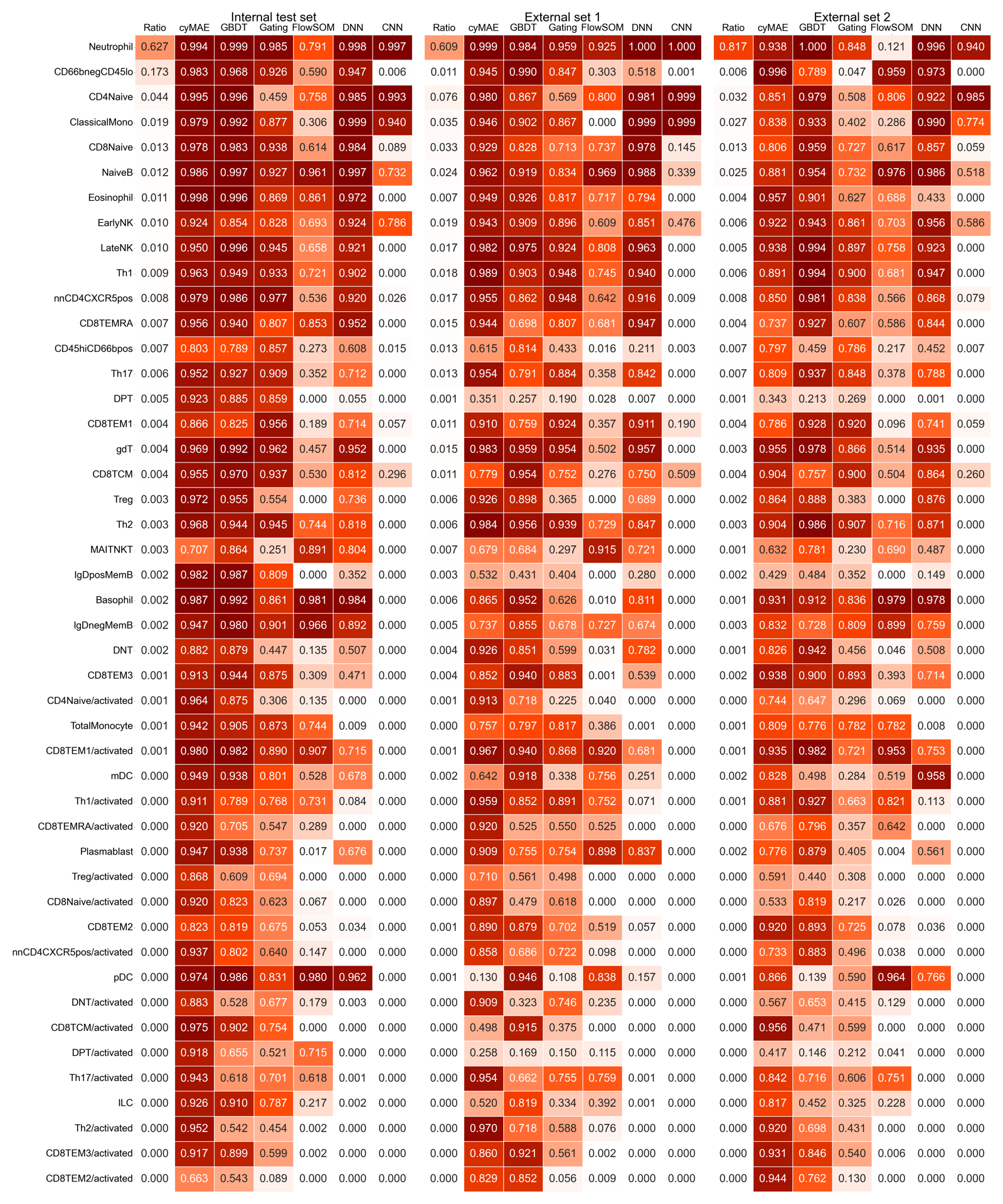

Figure S5. Full results of cell type prediction

Figure S6. The true expression and imputed expression plots for the Acute2020 dataset

Figure S7. The true expression and imputed expression plots for the Vaccine dataset.

Figure S8. The true expression and imputed expression plots for the Acute2021 dataset

.

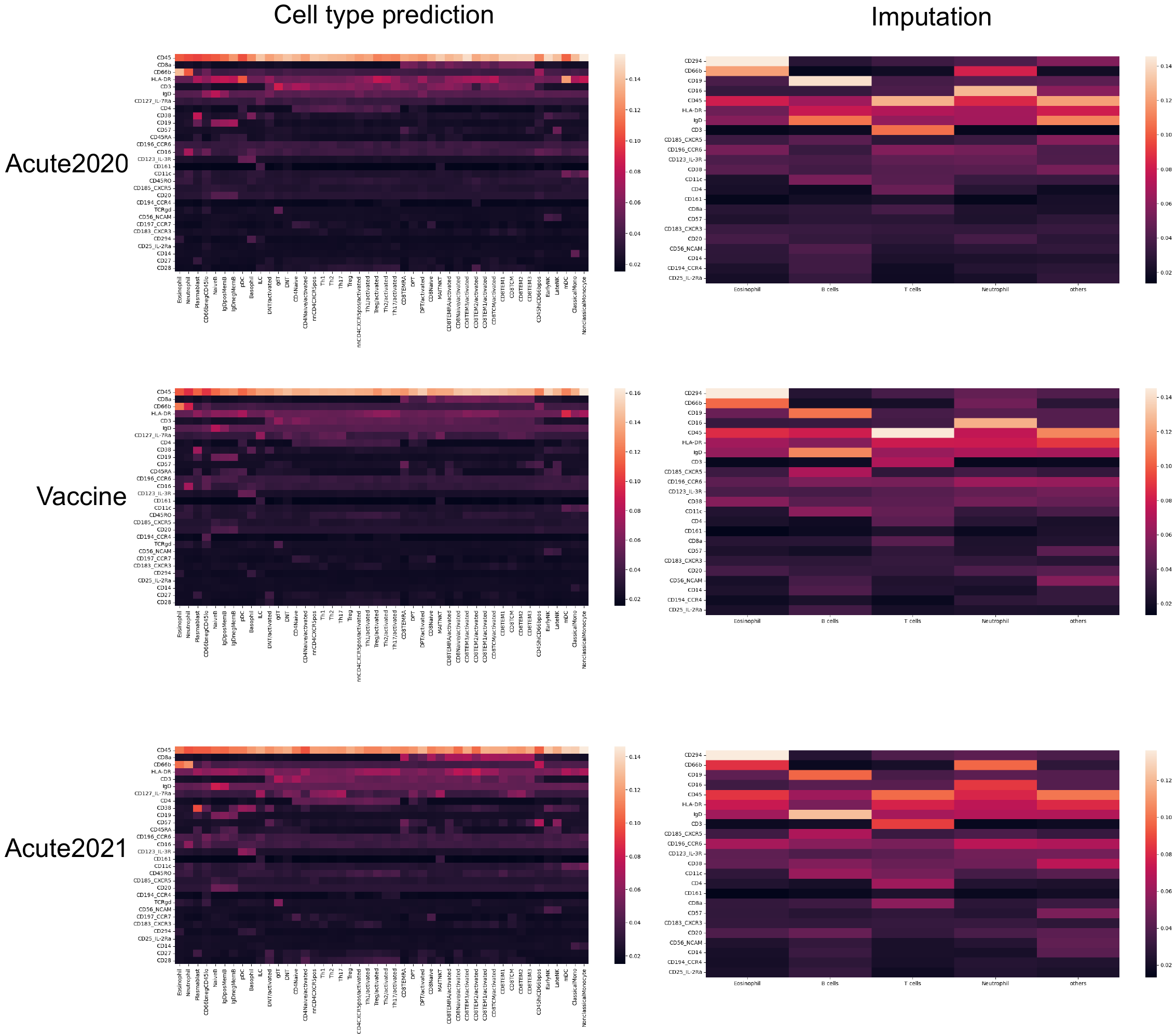

Figure S9. Attention scores of each cell type in the cell type classification and the imputation task.

Figure S10. Attention scores of each cell type for entire samples in the cell type classification task.

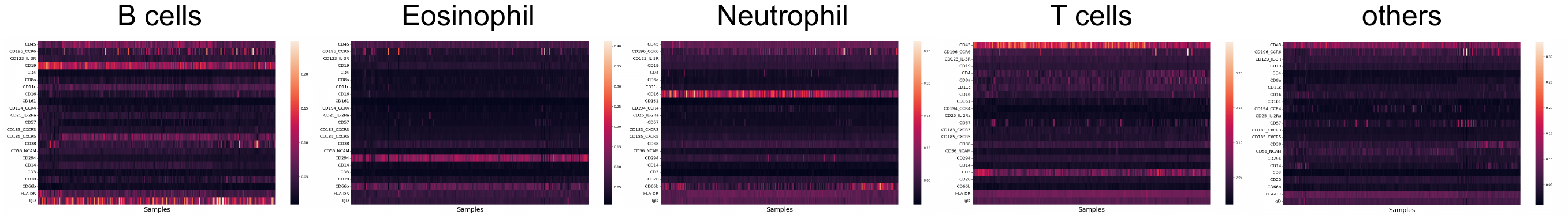

Figure S11. Attention scores of each cell type for entire samples in the imputation task.

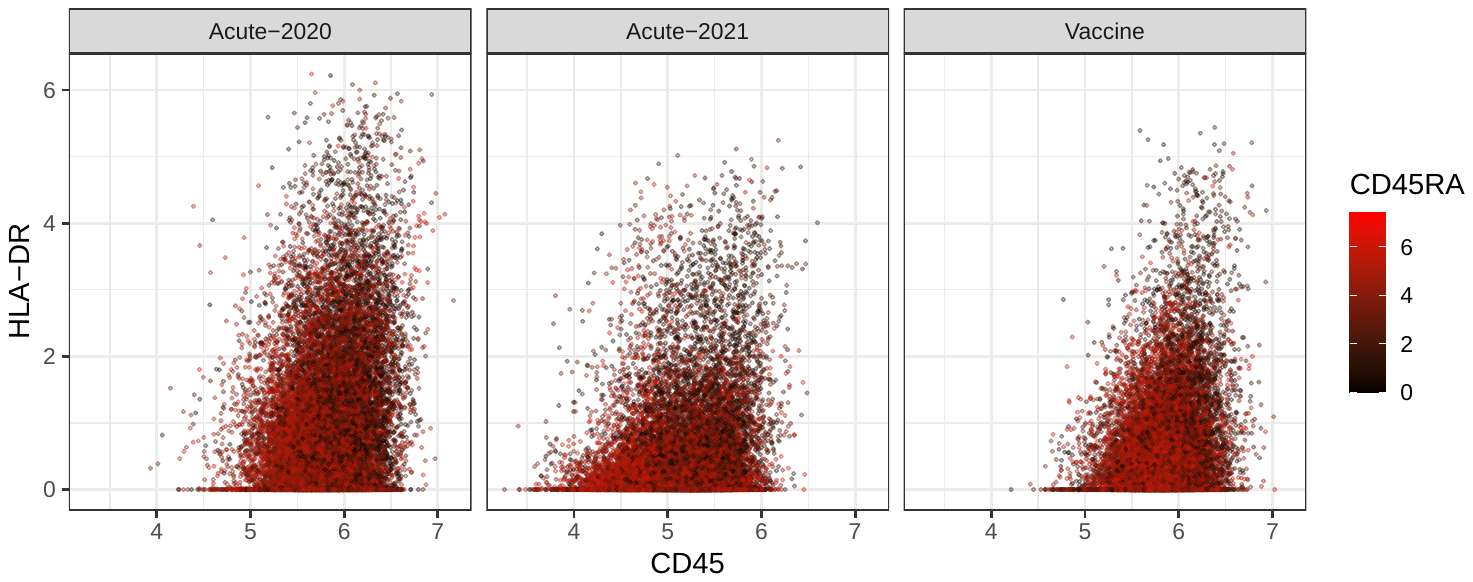

Figure S12. CD45 and HLA-DR both are negatively correlated to CD45RA in T cells

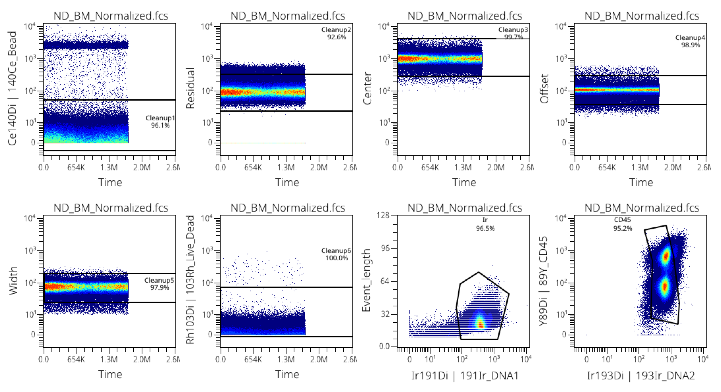

Figure S13. A standard cleanup procedure, which is a routine manual gating practice. fcs files were gated for beads, debris, doublets, and dead cells using the OMIQ platform.

Table S1. Balanced accuracy comparison between the non-pre-trained and the pre-trained cyMAE in cell type annotation.

|  | **Internal test set (Bacc)** | **External set 1 (Bacc)** | **External set2 (Bacc)** |
| --- | --- | --- | --- |
| cyMAE from scratch | 0.925 | 0.822 | 0.810 |
| cyMAE with fine-tuning | 0.931 | 0.825 | 0.810 |

Table S2. Full results of COVID-19 and healthy classification problem.

| **Feature extraction methods** | | **Predictors** | **Validation set**  **(AUROC Mean ± Std.)** | **Test set**  **(AUROC Mean ± Std.)** |
| --- | --- | --- | --- | --- |
| Manual gating | | GBDT | **0.970 ± 0.091** | **0.975 ± 0.042** |
|  |  | Logistic regression with L2 reg. | 0.917 ± 0.047 | 0.938 ± 0.059 |
| FlowSOM | | GBDT | **0.930 ± 0.089** | **0.936 ± 0.096** |
|  |  | Logistic regression with L2 reg. | 0.875 ± 0.077 | 0.902 ± 0.090 |
| cyMAE | Global mean pooling | Logistic regression with L2 reg. | 0.926 ± 0.068 | 0.930 ± 0.068 |
|  | Global sum pooling | Logistic regression with L2 reg. | 0.933 ± 0.068 | 0.922 ± 0.070 |
|  | Global max pooling | Logistic regression with L2 reg. | 0.989 ± 0.025 | 0.986 ± 0.035 |
|  | Global min pooling | Logistic regression with L2 reg. | **0.992 ± 0.021** | **0.987 ± 0.035** |

* reg. stands for regularization, Std. stands for standard deviation, and AUROC stands for Area Under the Receiver Operating Characteristic curve.

Table S3. Full results of Secondary immune response against COVID-19 prediction problem.

| **Feature extraction methods** | | **Predictors** | **Validation set**  **(AUROC Mean ± Std.)** | **Test set**  **(AUROC Mean ± Std.)** |
| --- | --- | --- | --- | --- |
| Manual gating | | GBDT | **0.735 ± 0.129** | **0.641 ± 0.154** |
|  |  | Logistic regression with L2 reg. | 0.456 ± 0.171 | 0.446 ± 0.140 |
| FlowSOM | | GBDT | **0.622 ± 0.132** | **0.579 ± 0.151** |
|  |  | Logistic regression with L2 reg. | 0.577 ± 0.162 | 0.520 ± 0.167 |
| cyMAE | Global mean pooling | Logistic regression with L2 reg. | 0.557 ± 0.133 | 0.561 ± 0.156 |
|  | Global sum pooling | Logistic regression with L2 reg. | 0.553 ± 0.119 | 0.599 ± 0.157 |
|  | Global max pooling | Logistic regression with L2 reg. | 0.671 ± 0.121 | 0.646 ± 0.147 |
|  | Global min pooling | Logistic regression with L2 reg. | **0.679 ± 0.119** | **0.669 ± 0.152** |

* reg. stands for regularization, Std. stands for standard deviation, and AUROC stands for Area Under the Receiver Operating Characteristic curve.

Table S4. Full results of COVID-19 pre- and post-treatment classification problem.

| **Feature extraction methods** | | **Predictors** | **Validation set**  **(AUROC Mean ± Std.)** | **Test set**  **(AUROC Mean ± Std.)** |
| --- | --- | --- | --- | --- |
| Manual gating | | GBDT | **0.865 ± 0.126** | **0.796 ± 0.124** |
|  |  | Logistic regression with L2 reg. | 0.621 ± 0.120 | 0.615 ± 0.177 |
| FlowSOM | | GBDT | **0.885 ± 0.106** | **0.859 ± 0.112** |
|  |  | Logistic regression with L2 reg. | 0.592 ± 0.134 | 0.579 ± 0.170 |
| cyMAE | Global mean pooling | Logistic regression with L2 reg. | 0.757 ± 0.200 | 0.708 ± 0.138 |
|  | Global sum pooling | Logistic regression with L2 reg. | 0.739 ± 0.164 | 0.703 ± 0.149 |
|  | Global max pooling | Logistic regression with L2 reg. | **0.921 ± 0.090** | **0.861 ± 0.104** |
|  | Global min pooling | Logistic regression with L2 reg. | 0.845 ± 0.133 | 0.806 ± 0.128 |

* reg. stands for regularization, Std. stands for standard deviation, and AUROC stands for Area Under the Receiver Operating Characteristic curve.

Table S5. Cell type annotation using different masking ratios in the cyMAE.

| **Masking ratio** | **Internal test set (Bacc)** | **External set 1 (Bacc)** | **External set 2 (Bacc)** |
| --- | --- | --- | --- |
| 0.25 | 0.931 | 0.825 | 0.810 |
| 0.5 | 0.931 | 0.824 | 0.813 |
| 0.75 | 0.932 | 0.825 | 0.812 |
